# An ancient deletion in the ABO gene affects the composition of the porcine microbiome by altering intestinal N-acetyl-galactosamine concentrations

**DOI:** 10.1101/2020.07.16.206219

**Authors:** Hui Yang, Jinyuan Wu, Xiaochang Huang, Yunyan Zhou, Yifeng Zhang, Min Liu, Qin Liu, Shanlin Ke, Maozhang He, Hao Fu, Shaoming Fang, Xinwei Xiong, Hui Jiang, Zhe Chen, Zhongzi Wu, Huanfa Gong, Xinkai Tong, Yizhong Huang, Junwu Ma, Jun Gao, Carole Charlier, Wouter Coppieters, Lev Shagam, Zhiyan Zhang, Huashui Ai, Bin Yang, Michel Georges, Congying Chen, Lusheng Huang

## Abstract

We have generated a large heterogenous stock population by intercrossing eight divergent pig breeds for multiple generations. We have analyzed the composition of the intestinal microbiota at different ages and anatomical locations in > 1,000 6^th^- and 7^th^- generation animals. We show that, under conditions of exacerbated genetic yet controlled environmental variability, microbiota composition and abundance of specific taxa (including *Christensenellaceae*) are heritable in this monogastric omnivore. We fine-map a QTL with major effect on the abundance of *Erysipelotrichaceae* to chromosome 1q and show that it is caused by a common 2.3-Kb deletion inactivating the ABO acetyl-galactosaminyl-transferase gene. We show that this deletion is a trans-species polymorphism that is ≥3.5 million years old and under balancing selection. We demonstrate that it acts by decreasing the concentrations of N-acetyl-galactosamine in the cecum thereby reducing the abundance of *Erysipelotrichaceae* strains that have the capacity to import and catabolize N-acetyl-galactosamine.

## Introduction

It is increasingly recognized that a comprehensive understanding of the physiology and pathology of organisms (including humans) requires the integrated analysis of the host and its multiple microbiota, i.e. to consider the organism as a “holobiont” (Kundu et al., 2017). In human, intestinal microbiota composition is significantly associated with physiological and pathological parameters including HDL cholesterol, fasting glucose levels and body mass index (BMI)(Rothschild et al., 2018). In livestock, ruminal microbiome composition is associated with economically important traits including methane production and feed efficiency (O’Hara et al., 2020). These correlations reflect a complex interplay between host and microbiota, which may include direct (“causal”) effects of the microbiome on the host’s physiology. This is supported by conventionalization experiments (aka human microbiota-associated (HMA) rodents), although it has been rightfully pointed out that the conclusions of many of these experiments have to be considered with caution (Walter et al., 2020). Several of the phenotypes correlated with microbiota composition, whether in humans or animals, are heritable in the sense that a significant fraction of the trait variance can be explained by genetic differences between individuals (Falconer & Mackay, 1996; Polderman et al., 2015; Polubriaginof et al., 2018). Combined, this leads to the intriguing hypothesis that the genotype of the host may determine the composition of the microbiota which may in turn affect the host’s phenotype (Goodrich et al., 2014; Schmidt et al., 2018). This assertion implies that the composition of the microbiota is itself heritable. While mapping data in rodents support this hypothesis (Schlamp et al., 2019), the evidence has been shallower in humans. Initial reports didn’t reveal a higher microbiota resemblance between monozygotic than dizygotic twins suggesting limited impact of host genotype on microbiota composition (Yatsunenko et al., 2012). Yet better-powered studies using larger twin cohorts provided evidence for a significant impact of host genetics on the abundance of some taxa, particularly the family *Christensenellaceae* (Goodrich et al., 2014). Specific loci that may underpin microbiota heritability have remained difficult to identify. Apart from chromosome 2 variants that cause persistent expression of lactase (*LCT*) in the gut which have reproducibly been found associated with increased proportions of *Bifidobacterium* (probably as a consequence of altered dietary habits due to lactose tolerance), other GWAS-identified loci have proven more difficult to replicate (Blekhman et al., 2015, Turpin et al., 2016, Bonder et al., 2016, Wang et al., 2016, Rothschild et al., 2018, Hughes et al., 2020). The analysis of larger human cohorts is needed to gain a better understanding of the likely highly polygenic genetic architecture of microbiota composition.

In an effort to contribute to deciphering the genetic architecture of intestinal microbiota composition in an omnivorous, monogastric model of size comparable to human we herein report the generation of a mosaic pig population by intercrossing eight divergent founder breeds for up to seven generations (hence exacerbating genetic variation), and the longitudinal characterization of the intestinal microbiome of F6 and F7 animals that were raised in uniform conditions (hence minimizing environmental variation). We provide evidence for a strong impact of host genotype on microbiome composition and identify a locus with large effect on the abundance of specific taxa by controlling the concentration of a particular metabolite in the gut thereby affecting species that can use this metabolite as carbon source.

## Results

### Generating a large mosaic pig population for genetic analysis of complex phenotypes

We have generated a large (> 7,500 animals in total) multigenerational (> seven generations) heterogeneous or mosaic population by inter-crossing the offspring of 61 founder animals (F0) representing four aboriginal Chinese breeds and four commercial European and American breeds using a rotational design (Fig. 1A; Suppl. Table 1). All animals were reared in standardized housing and feeding conditions at one location (see Methods). We measured >200 phenotypes (pertaining to body composition, physiology, disease resistance and behavior), transcriptome, epigenome and chromatin interaction data on multiple tissues, plasma metabolome and microbiome data (see hereafter) in up to 954 F6 and 892 F7 animals. All F0 animals were whole-genome sequenced at average 28.4-fold depth (range: 23.1 – 37.2), while all F6 and F7 animals were sequenced at average 8.0-fold depth (range: 5.2-12.4). SNPs were detected and genotypes called using Platypus yielding 23.8 million SNPs and 6.4 million indels with MAF ≥ 0.03 (in F0, F6 and F7 combined) for further analyses. The nucleotide diversity averaged 2.5×10^−3^ within Chinese founders, 2.0×10^−3^ within European founders, 3.6×10^−3^ between Chinese founders, 2.5×10^−3^ between European founders and 4.3×10^−3^ between Chinese and European founders (Fig. 1B). We used a linear model incorporating all variants to estimate the average contribution of the eight founder breeds in the F6 and F7 generation at genome and chromosome level (Coppieters et al., 2020). At genome-wide level, the proportion of the eight founder breed genomes ranged from 11.2% (respectively 11.5%) to 14.1% (14.7%) in the F6 (F7) generations. At chromosome-specific level, the proportion of the eight founder breeds ranged from 6.7% (respectively 4.9%) to 20.7% (22.1%) in the F6 (F7) generations (Fig. 1C). As indicators of mapping resolution achievable in this cross, the median number of variants in high linkage disequilibrium (LD) (*r*^2^ ≥ 0.9) with a reference variant was 30 (Fig. 1D), and the median maximal distance with a high variant in high LD (*r*^2^ ≥ 0.9) was 54,015 base pairs (Fig. 1E).

**Figure 1:**
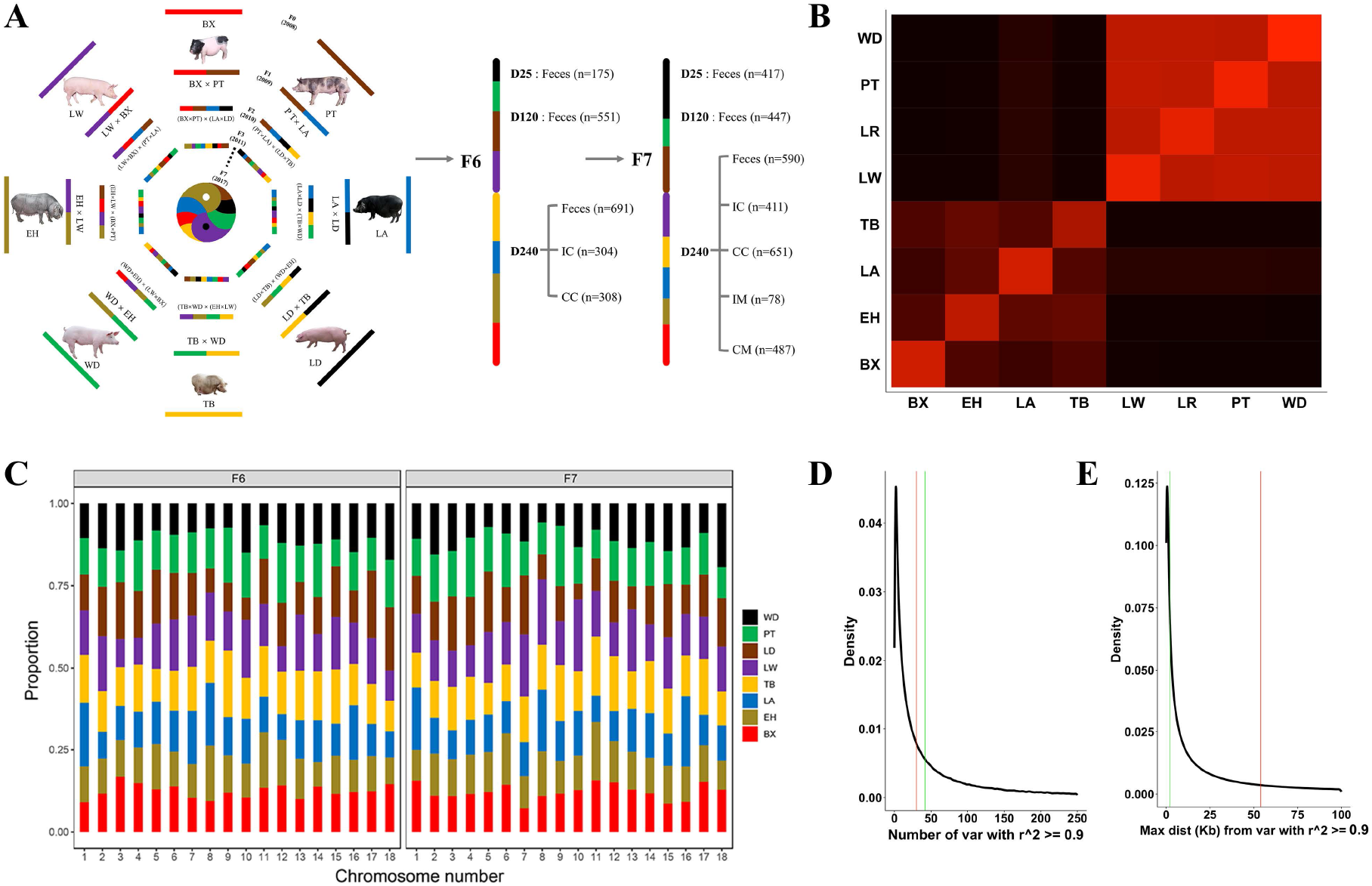
**(A)** Rotational breeding design used for the generation of a large mosaic pig population for the genetic analysis of complex phenotypes, with sampling scheme for feces (D25, D120, D240), luminal content of the ileum (IC) and cecum (CC), and mucosal scrapings in the ileum (IM) and cecum (CM). BX: Bamaxiang, EH: Erhualian, LA: Laiwu, TB: Tibetan, LW: Large White, LD: Landrace, PT: Piétrain, WD: White Duroc. **(B)** Average similarity (1 – *π*) between allelic sequences sampled within and between the eight founder breeds. The color intensity ranges from black (breeds with lowest allelic similarity: BX vs WD, 1-4.3×10^−3^) to bright red (breed with highest allelic similarity: WD, 1-1.8×10^−3^). The acronyms for the breeds are as in (A). Within-breed similarity is higher than between-breed similarity as expected. Between-breed similarity is lower for Chinese than for European breeds, and still lower between Chinese and European breeds. Laiwu are slightly more similar to European breeds than the other Chinese breeds. **(C)** Autosome-specific estimates of the genomic contributions of the eight founder breeds in the F6 and F7 generation. **(D)** Frequency distribution (density) of the number of variants in high LD (*r*^2^ ≥ 0.9) with a reference variant, corresponding to the expected size of “credible sets” in GWAS (Huang et al., 2017). The red vertical line corresponds to the genome-wide median. The green vertical line corresponds to the mapping resolution achieved in this study for the ABO locus (see hereafter). **(E)** Frequency distribution (density) of the maximum distance between a reference variant and a variant in high LD (*r*^2^ ≥ 0.9) with it defining the spread of credible sets. Red and green vertical lines are as in (D).

**Supplemental Table 1:** Numbers of parents used and animals produced for the different generations of the mosaic pig population.

| Generation | Total nr animals produced | Nr animals used as boars | Nr animals used as sows |
| --- | --- | --- | --- |
| F0 | 61 | 29 | 32 |
| F1 | 265 | 32 | 58 |
| F2 | 575 | 56 | 87 |
| F3 | 776 | 57 | 75 |
| F4 | 746 | 62 | 97 |
| F5 | 938 | 85 | 170 |
| F6 | 1663 | 82 | 111 |
| F7 | 1227 | 72 | 94 |
| F8 | 780 | 66 | 83 |
| F9 | 595 | 62 | 56 |

**Supplemental Table 2:** 16S rRNA based microbiome profiling of 12 data sets: summary statistics

| Generation | Dataset | Sample type | Sample size | Number of tags |  |  | Number of OTUs (selected) |  |  |  | Number of Taxa |  |  |  |  |  |
| --- | --- | --- | --- | --- | --- | --- | --- | --- | --- | --- | --- | --- | --- | --- | --- | --- |
|  |  |  |  | Mean | Min | Max | Mean | Min | Max | Total | Phylum | Class | Order | Family | Genus | Species |
| F6 | D25 | Feces | 175 | 34,153 | 25,486 | 43,557 | 804 | 229 | 1,640 | 8,085 | 34 | 77 | 130 | 182 | 300 | 135 |
|  | D120 | Feces | 551 | 33,378 | 22,807 | 43,447 | 1,751 | 651 | 2,616 | 10,657 | 31 | 72 | 121 | 171 | 285 | 123 |
|  | D240 | Feces | 691 | 32,024 | 19,632 | 43,045 | 2,048 | 1,168 | 2,884 | 11,291 | 40 | 83 | 144 | 197 | 339 | 140 |
|  | IC | Ileal content | 304 | 34,034 | 24,993 | 43,854 | 378 | 60 | 2,333 | 6,448 | 40 | 85 | 145 | 200 | 341 | 143 |
|  | CC | Cecum content | 308 | 34,005 | 23,593 | 43,674 | 1,446 | 622 | 2,663 | 9,876 | 25 | 50 | 92 | 145 | 249 | 119 |
| F7 | D25 | Feces | 417 | 45,094 | 20,993 | 79,771 | 978 | 264 | 2,115 | 9,738 | 24 | 42 | 72 | 122 | 235 | 118 |
|  | D120 | Feces | 447 | 41,796 | 26,425 | 65,235 | 2,285 | 491 | 3,005 | 10,362 | 23 | 39 | 64 | 110 | 195 | 94 |
|  | D240 | Feces | 590 | 40,915 | 24,010 | 61,986 | 2,422 | 1,138 | 3,112 | 10,226 | 23 | 41 | 62 | 101 | 189 | 96 |
|  | IC | Ileal content | 411 | 54,700 | 29,825 | 73,709 | 241 | 54 | 1,120 | 4,773 | 30 | 64 | 119 | 170 | 304 | 132 |
|  | IM | Ileal mucosa | 78 | 52,239 | 29,860 | 73,685 | 1,024 | 217 | 2,014 | 7,722 | 30 | 64 | 116 | 165 | 298 | 131 |
|  | CC | Cecum content | 651 | 47,796 | 30,835 | 62,424 | 2,083 | 295 | 2,892 | 10,416 | 22 | 38 | 66 | 113 | 220 | 115 |
|  | CM | Cecum mucosa | 487 | 45,836 | 27,520 | 71,481 | 1,179 | 279 | 2,243 | 10,265 | 27 | 58 | 110 | 161 | 289 | 132 |

### Characterizing the age- and location-specific composition of the intestinal microbiome of the healthy pig

We collected feces at 25 days (i.e. suckling period), 120 days (i.e. growing period) and 240 days (i.e. day of slaughter), as well as cecal and ileal content and mucosal scrapings (F7 only) at day 240 in the F6 and F7 generations (total of 7 traits and 12 data series) (Fig. 1A). We performed 16S rRNA sequencing (V3-V4 hypervariable region) and obtained usable post-QC data for 5,110 samples. Sample size per data series averaged 426 (range: 78-691) (Suppl. Table 2). Sequence tags were rarefied to ~20,000 per sample, and clustered in 32,032 OTUs (97% similarity threshold). 12,054 OTUs present in at least 0.2% of the samples (with more than two tags in at least two samples) and amounting to an average of 98.7% of sample reads, were retained for further analysis. They were annotated to 41 phyla, 87 classes, 149 orders, 207 families, 360 genera and 150 species. The number of OTUs detected per sample averaged 1,575 (range: 54 to 3,112) (Suppl. Table 2). The first two Principal Coordinates (based on Bray-Curtis distance) separated the samples by trait consistently across the two cohorts, generating five dominant clusters: (i) day 25 feces, (ii) day 120 and 240 feces, (iii) cecal content and mucosa, (iv) ileal content, and (v) ileal mucosa (Fig. 2A). Fecal samples were dominated by *Firmicutes* and *Bacteroidetes*. Day 25 samples had larger proportions of *Proteobacteria* and *Fusobacteria*, while day 120 and day 240 samples had larger proportions of *Spirochaetes*. Cecum content and mucosa had lower proportion of *Firmicutes* and *Spirochaetes* than day 120-240 feces, yet higher proportions of *Bacteroidetes, Proteobacteria* and *Fusobacteria*. Ileal samples differed more dramatically from the others. They had much lower proportions of *Bacteroidetes*, were dominated by *Clostridiaceae* (= *Firmicutes*) and *Enterobacteriaceae* (= *Proteobacteria*), and had a high proportion of *Pasteurellaceae* (= *Proteobacteria*). Ileal mucosa differed considerably from ileal content, having a higher proportion of *Bacteroidetes* and *Spirochaetes*, yet less *Firmicutes* and *Proteobacteria* (Fig. 2B and Suppl. Table 3). A total of 58 OTUs that were annotated to 21 taxa were identified in >95% of day 120 and 240 feces and cecum content samples of both F6 and F7 generations, hence defined as core bacterial taxa (Suppl. Fig. 1A). *α*-diversity (measured by Shannon’s index) of fecal samples was lower at day 25 than at days 120-240, reminiscent of the enrichment of the intestinal flora observed between child- and adult-hood in humans (Yatsunenko et al., 2012; Radjabzadeh et al., 2020). It was also lower for ileal content than for cecal content and mucosa (and probably ileal mucosa) (Fig. 2C). Six percent of ileal content samples harbored less than 100 OTUs (Suppl. Table 2). *β*-diversity (measured by pair-wise Bray-Curtis dissimilarities) tended to be inversely proportional to *α*-diversity, being higher for day 25 than for day 120 and 240 fecal samples. The variation of pair-wise Bray-Curtis dissimilarities was highest for ileal content which had the lowest *α*-diversity (Fig. 2D). Microbiota composition of pig D240 feces was more similar to that of human than of mice feces (Suppl. Fig. 1B). Human feces contained more *Proteobacteria* and less *Spirochaetes*, while mice feces contained more *Firmicutes* yet less *Bacteroidetes* and *Spirochaetes* (Fig. 2B)

**Figure 2:**
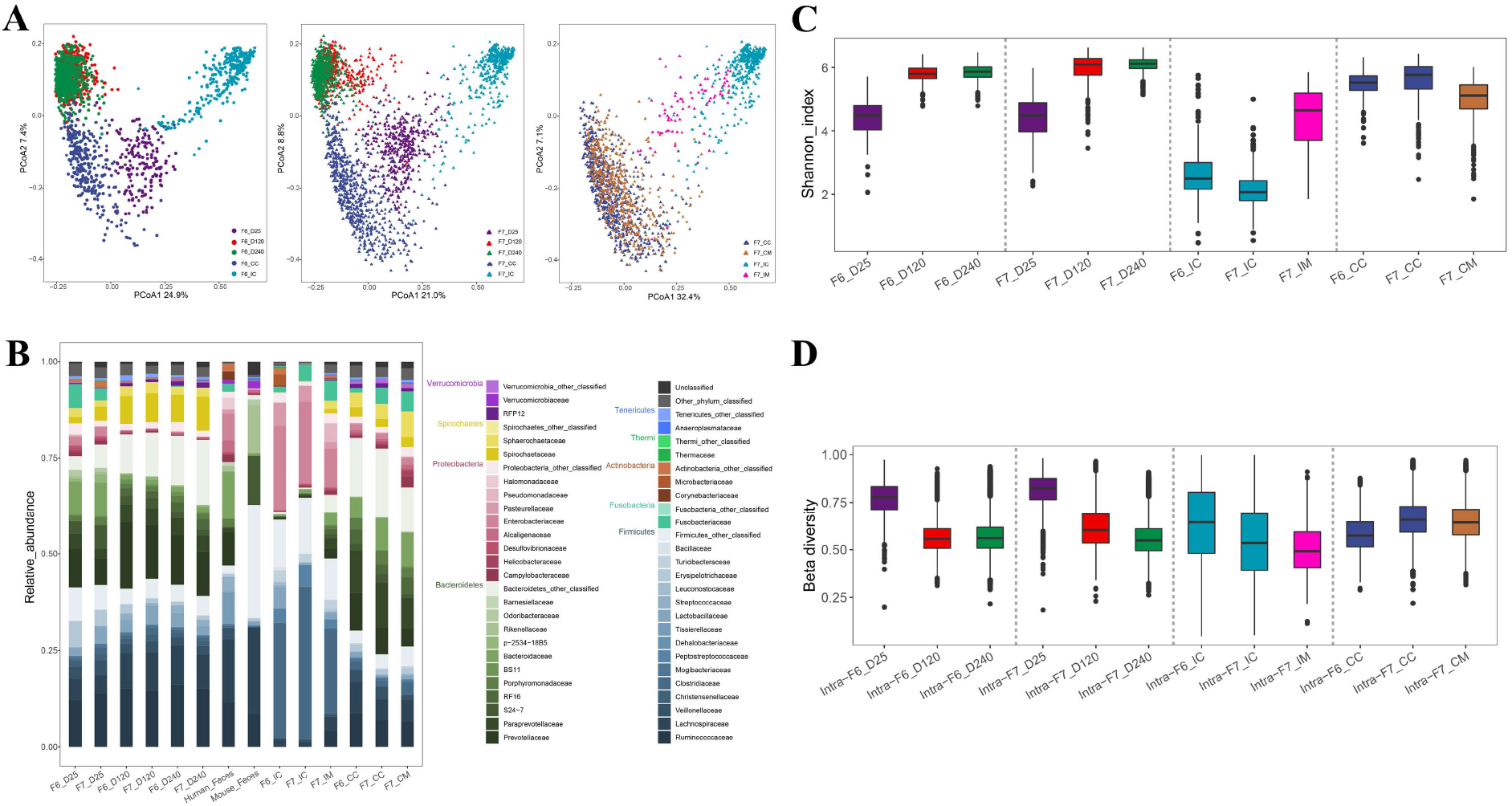
**(A)** Joint Principal Coordinate Analysis (PCoA) of 5,110 16S rRNA microbiome profiles. (I) Generation F6: fecal samples day 25 (D25, mauve), fecal samples day 120 (D120, red), fecal samples day 240 (D240, green), ileal content (IC, light blue), cecum content (CC, dark blue). (II) Generation F7: fecal samples day 25 (D25, mauve), fecal samples day 120 (D120, red), fecal samples day 240 (D240, green), ileal content (IC, light blue), cecum content (CC, dark blue). (III) Generation F7: ileal content (IC, light blue), cecum content (CC, dark blue), ileal mucosa (IM, pink), cecal mucosa (CM, brown). **(B)** Average 16S rRNA microbiota composition of the 12 porcine data series. Taxa are colored by phylum and by family within phylum, highlighting 43 families that are amongst the top 15 in at least one data series. The names of the corresponding phyla and families are provided in the legend. Average microbiota composition of 106 human feces and 6 mouse feces (C57BL/6). **(C)** Individual *α*-diversities measured using Shannon’s index for the 12 porcine data series color-labelled as in A and B. **(D)** Individual *β*-diversities measured pair-wise Bray-Curtis dissimilarities for the 12 porcine data series color-labelled as in A and B.

**Suppl. Fig. 1:**
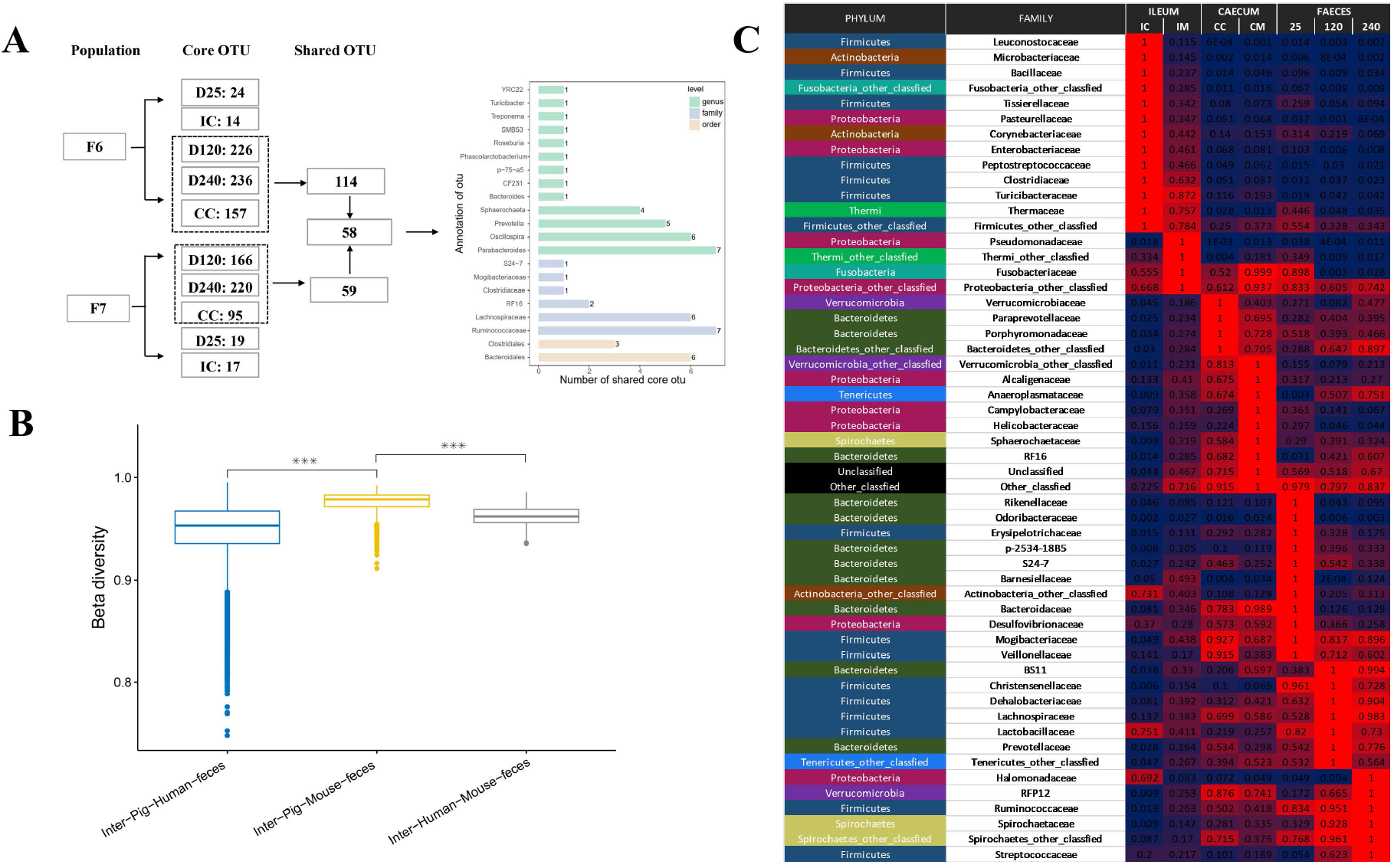
**(A)** Definition of a core intestinal microbiome of the pig. **(B)** The compositions of the porcine and human intestinal microbiota are closer to each other than either is to that of the mouse. **(C)** Abundances (F6-F7 averages when available) of the 43 families represented in Fig. 2B in the seven types of samples (“traits”) relative to the sample type in which they are the most abundant (red – blue scale). The families are ordered according to the sample type in which they are the most abundant. The color-code for phyla is as in Fig. 2B.

**Supplemental Table 3:** 16S rRNA (V3-V4) based abundances of 43 bacterial families that are amongst the top 15 in at least one of the 12 data-series (used for figure 2B).

| Taxon | Phylum | Family | Average abundance | Data series |
| --- | --- | --- | --- | --- |
| Bacteria;Bacteroidetes;Bacteroidia;Bacteroidales;Paraprevotellaceae | Bacteroidetes | Paraprevotellaceae | 0.109390596 | F6_CC |
| Bacteria;Bacteroidetes;Bacteroidia;Bacteroidales;Prevotellaceae | Bacteroidetes | Prevotellaceae | 0.097737987 | F6_CC |
| Bacteria;Firmicutes;Clostridia;Clostridiales;Ruminococcaceae | Firmicutes | Ruminococcaceae | 0.087122887 | F6_CC |
| Bacteria;Firmicutes;Clostridia;Clostridiales;Lachnospiraceae | Firmicutes | Lachnospiraceae | 0.08153471 | F6_CC |
| Bacteria;Bacteroidetes;Bacteroidia;Bacteroidales;Bacteroidaceae | Bacteroidetes | Bacteroidaceae | 0.053424685 | F6_CC |
| Bacteria;Spirochaetes;Spirochaetes;Sphaerochaetales;Sphaerochaetaceae | Spirochaetes | Sphaerochaetaceae | 0.038308145 | F6_CC |
| Bacteria;Bacteroidetes;Bacteroidia;Bacteroidales;RF16 | Bacteroidetes | RF16 | 0.03501472 | F6_CC |
| Bacteria;Firmicutes;Clostridia;Clostridiales;Veillonellaceae | Firmicutes | Veillonellaceae | 0.031823326 | F6_CC |
| Bacteria;Bacteroidetes;Bacteroidia;Bacteroidales;Porphyromonadaceae | Bacteroidetes | Porphyromonadaceae | 0.03073182 | F6_CC |
| Bacteria;Spirochaetes;Spirochaetes;Spirochaetales;Spirochaetaceae | Spirochaetes | Spirochaetaceae | 0.024133387 | F6_CC |
| Bacteria;Firmicutes;Clostridia;Clostridiales;Clostridiaceae | Firmicutes | Clostridiaceae | 0.020539321 | F6_CC |
| Bacteria;Bacteroidetes;Bacteroidia;Bacteroidales;S24-7 | Bacteroidetes | S24-7 | 0.019756632 | F6_CC |

(12 first rows only)

### Evaluating the heritability of intestinal microbiome composition in the mosaic pig population

To evaluate to what extent individual genotype contributes to the observed *β*-diversity (i.e. measure the heritability of microbiota composition), we regressed pair-wise Bray-Curtis dissimilarity on pair-wise kinship coefficient measured from genome-wide SNP data (Rothschild et al., 2018). We first performed analyses within litter following Visscher et al. (2006). There is no reason to assume that litter-mates that are genetically more similar to each other would also be exposed to more similar environmental effects. Hence, a significant negative correlation between genetic similarity and microbiome dissimilarity within litter is strong evidence for an effect of genetics on microbiome composition. Correlations were measured separately for the 12 measured data series (day 25, day 120, day 240 fecal samples (F6 & F7), ileal and cecal content (F6 & F7), and ileal and cecal mucosae (F7)). The number of litters per analysis averaged 90 (range: 18 to 156), while the number of full-sib pairs per analysis averaged 587 (range: 109 – 1,215). The range of kinship and Bray-Curtis dissimilarity values may differ between litters, whether by chance, as a result of idiosyncrasies of the parental SNP genotypes, and/or of differences in environmental conditions between litters, and this may inflate the correlations (both up and down-wards). Thus, we evaluated the statistical significance of the observed correlations by performing 1,000 permutations of kinship coefficients and Bray-Curtis dissimilarities within litter. The correlations were negative for the 12 analyzed traits, and below the 50ties percentile of the permutation values for 11 of 12 (p = 0.0029) (not in IM which has one of the smallest n). The empirical p-value (one-sided) of the correlation was ≤ 0.05/12=0.004 (Bonferroni corrected threshold) for two (D240 in F6 and F7, which have large n). We combined the p-values across the 12 data series by summing the ranks of the observed correlations and computing the probability of this sum (one-sided) under the null hypothesis by simulation (see Methods) yielding an overall p-value of 3 x 10^−4^, hence providing a first line of evidence for an effect of genetics on microbiome composition in this population (Fig. 3A).

**Figure 3:**
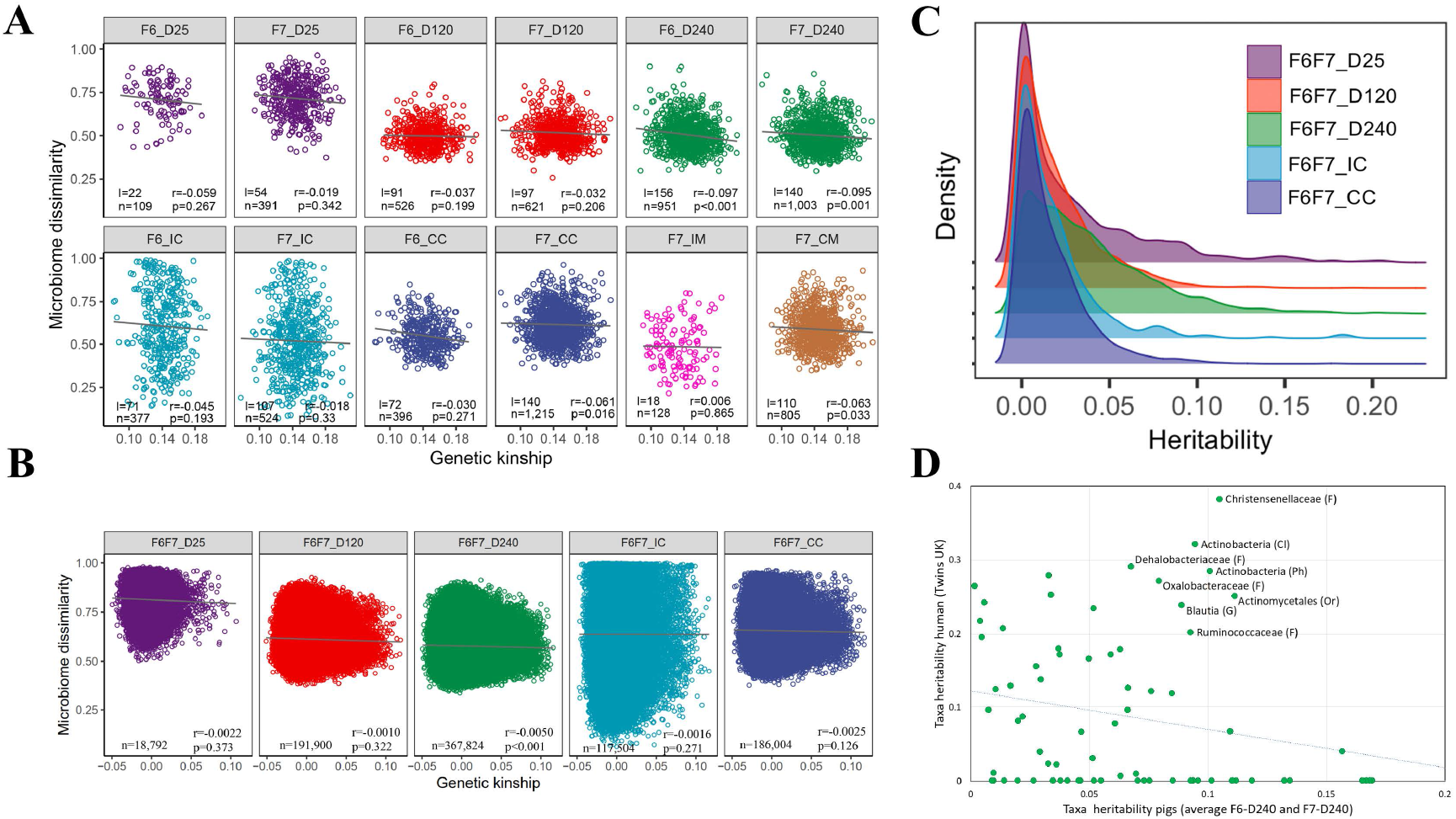
**(A) Correlation between genome-wide kinship (Θ) and microbiome dissimilarity (Bray-Curtis distance) within litter**. Correlations were measured separately for the 12 data series. P-values (one-sided p) were computed using a permutation test. Spearman’s correlations (r) were computed in R and adjusted to match the permutation p-values (see Methods). The number of litters (*l*) and animal pairs (*n*) used for analysis are given for each data series. **(B) Correlation between genome-wide kinship (Θ) and microbiome dissimilarity (Bray-Curtis distance) across generations**. We considered all possible pairs of F6 and F7 animals (ignoring sow-offspring pairs), hence considerably increasing sample size when compared to (A). Analyses were conducted for the five traits measured in both F6 and F7. *r, p*, and *n* are as in (A). **(C) Frequency distribution of heritabilities of individual taxa** for fecal samples (D25, D120 and D240) and intestinal content samples (IC and CC). Values are F6 and F7 averages. **(D) Correspondence between taxa heritabilities in human and pig adult fecal samples**.

We performed the same correlation analysis between genome-wide kinship and microbiome dissimilarity across the F6 and F7 generations (raised respectively in 2016 and 2017) for the five traits that were measured in both cohorts. None of the F6-F7 pairs considered included parent-offspring pairs (the microbiome of F6 sows may determine the microbiome of F7 offspring independently of genetics). The number of pairs in the across-generation analyses averaged 176,405 (range: 18,792 – 367,824). For the reasons described above, the statistical significance of the correlations was determined by 1,000 permutations performed within F6 and F7 litters (see Methods). The correlation was negative and below the 50-ties percentile of the permutation values for the five analyzed traits (p = 0.03). The empirical p-value (one-sided) of the correlation was ≤ 0.05/5=0.01 (Bonferroni corrected threshold) for one (D240 which has largest n). The combined p-value for the five traits combined and computed as above was 0.013, hence providing a second line of evidence for an effect of genetics on microbiome composition in this population (Fig. 3B).

We then evaluated the heritability (*h*^2^) of the abundances of individual taxa using a mixed model. This was done for up to 29 phyla, 53 classes, 86 orders, 116 families, 148 genera and 4,240 OTUs per data series. Heritabilities were estimated using a mixed model implemented with GEMMA (Zhou & Stephens, 2012). The model included random polygenic and residual error effects. Kinship coefficients (to constrain the polygenic effect) were computed from whole-genome SNP data, also using GEMMA (Zhou & Stephens, 2012). To obtain unbiased **h*^2^* estimates and associated p-values, we repeated the analysis 1,000 times with abundances randomly permuted within litter. The average *h*^2^ across the permutations was then subtracted from the *h*^2^ obtained with the unpermuted data to yield conservative estimates of 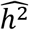. Their p-values were estimated as the proportion of permutations yielding an equally high or higher *h*^2^ estimate. Analyses were conducted for the 12 measured data series. P-values were ≤ 0.05 for 4,219 (=14%) of the 30,127 realized tests, hence above random expectations (Suppl. Table 4). The correlation between F6 and F7 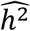 estimates (or their log(1/p) values) were positive and highly significant (p ≤ 1.1×10^−30^ and 1.07×10^−17^, respectively) for D240 fecal samples, hence supporting genuine genetic effects at least for this trait (Suppl. Fig. 2A). Averaged (over F6 and F7) heritabilities of individual taxa tended to be higher for fecal samples (especially at D240) than for content traits (Fig. 3C). Accordingly, total heritabilities computed following Rothschild et al. (2018) were highest for D240 fecal samples (5.09%) (Suppl. Fig. 2B). We compared the average heritabilities of individual taxa in D240 fecal samples with heritabilities of individual taxa in human feces (Goodrich et al., 2016). There was no significant correlation between pig and human values when considering all taxa. It is noteworthy, however, that the taxon found to be the most heritable in human, the family *Christensenellaceae* (Goodrich et al., 2014&2016), was also amongst the most heritable in pigs. At least seven other taxa were found to be heritable in both human and porcine adult feces (Fig. 3D).

**Suppl. Fig. 2:**
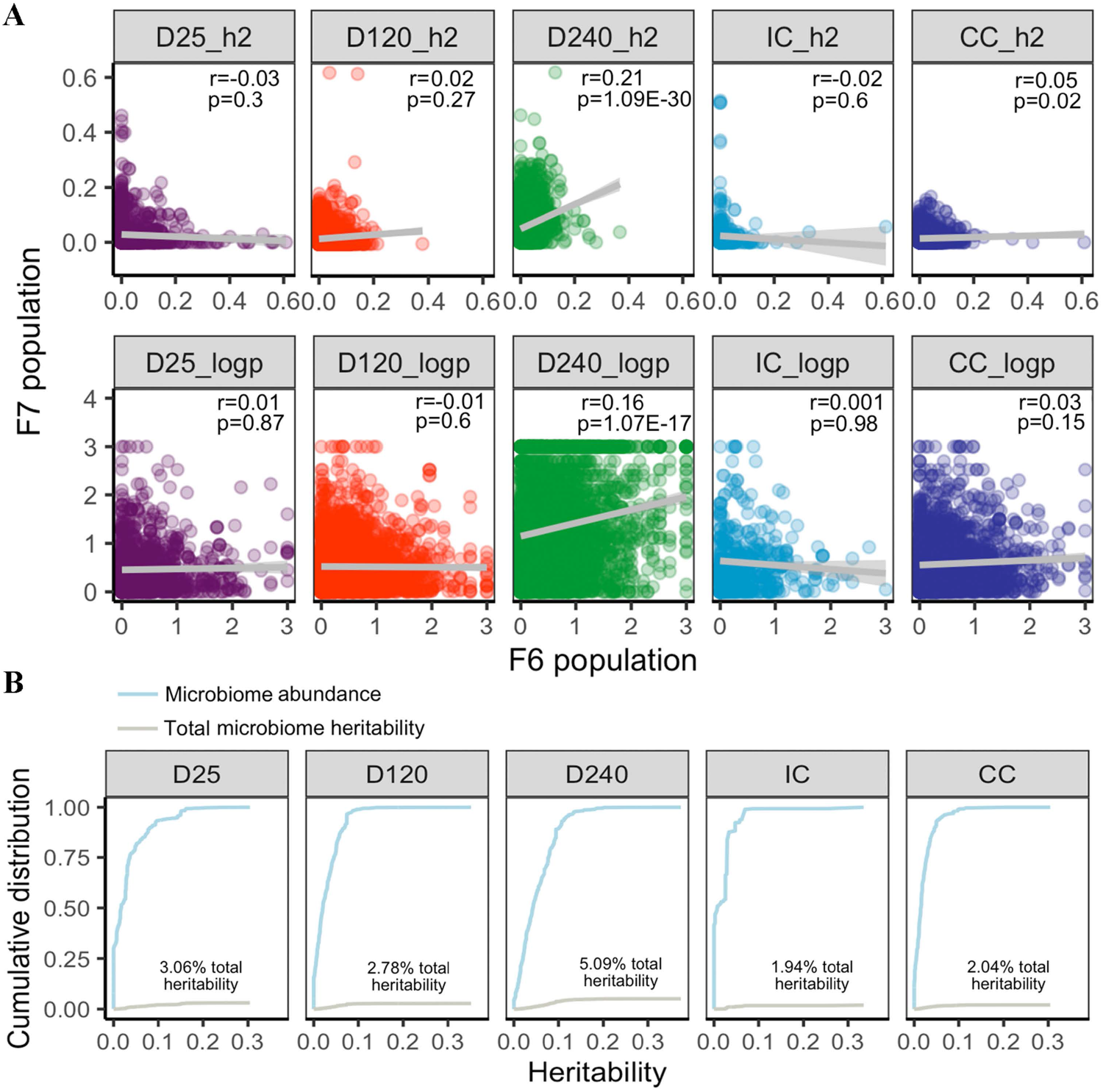
**(A)** Correlation between heritabilities (upper row) and associated log(1/p) values (lower row) of abundance of individual taxa between the F6 and F7 generations. Correlation coefficients (r) and corresponding p-values (p) are given. **(B)** Total heritabilities computed following Rothschild et al. (2018) using heritabilities of individual taxa averaged over the F6 and F7 generations for the five shared traits.

**Supplemental Table 4:** Heritabilities of indivdiual taxa for the 12 analyzed dataseries or traits.

| Data series / Trait | Taxon | Annotation | Uncorrected $h^2$ | Permutation $h^2$ | Corrected $h^2$ | p-value | q-value |
| --- | --- | --- | --- | --- | --- | --- | --- |
| F6_CC | Otu15777 | Bacteria;Firmicutes;Clostridia;Clostridiales;Ruminococcaceae;Ruminococcus | 0.155 | 0.043 | 0.112 | 0.00E+00 | 0.00E+00 |
| F6_CC | Otu28641 | Bacteria;Firmicutes;Clostridia;Clostridiales;Ruminococcaceae;Ruminococcus | 0.210 | 0.128 | 0.082 | 0.00E+00 | 0.00E+00 |
| F6_CC | Otu653 | Bacteria;Bacteroidetes;Bacteroidia;Bacteroidales | 0.084 | 0.004 | 0.080 | 0.00E+00 | 0.00E+00 |
| F6_CC | Otu654 | Bacteria;Bacteroidetes;Bacteroidia;Bacteroidales;Porphyromonadaceae | 0.130 | 0.047 | 0.083 | 0.00E+00 | 0.00E+00 |
| F6_CC | Otu7355 | Bacteria;Firmicutes;Clostridia;Clostridiales;Ruminococcaceae | 0.234 | 0.071 | 0.163 | 0.00E+00 | 0.00E+00 |
| F6_CC | Otu7838 | Bacteria;Firmicutes;Clostridia;Clostridiales;Ruminococcaceae;Oscillospira | 0.319 | 0.165 | 0.153 | 0.00E+00 | 0.00E+00 |
| F6_CC | Otu10305 | Bacteria;Bacteroidetes;Bacteroidia;Bacteroidales;Porphyromonadaceae;Parabacteroides | 0.157 | 0.047 | 0.110 | 1.00E-03 | 2.05E-01 |
| F6_CC | Otu16289 | Bacteria;Cyanobacteria;4C0d-2;YS2 | 0.240 | 0.145 | 0.095 | 1.00E-03 | 2.05E-01 |
| F6_CC | Otu2033 | Bacteria;Firmicutes;Clostridia;Clostridiales | 0.192 | 0.048 | 0.144 | 1.00E-03 | 2.05E-01 |
| F6_CC | Otu3738 | Bacteria;Spirochaetes;Spirochaetes | 0.286 | 0.121 | 0.165 | 1.00E-03 | 2.05E-01 |
| F6_CC | Otu6981 | Bacteria;Firmicutes;Clostridia;Clostridiales;Ruminococcaceae | 0.240 | 0.128 | 0.112 | 1.00E-03 | 2.05E-01 |
| F6_CC | Otu16266 | Bacteria;Cyanobacteria;4C0d-2;YS2 | 0.165 | 0.057 | 0.108 | 2.00E-03 | 3.01E-01 |

(12 first rows only)

### Identifying a microbiota QTL with major effect on the abundance of *Erysipelotrichaceae* species by whole genome sequence based GWAS

Having established that host genetics affects intestinal microbiome composition in our population, we sought to identify contributing loci by performing GWAS. GWAS were initially performed separately by trait, taxon and generation, and conducted using two statistical models following Turpin et al. (2016). The first model analyzed the effect of individual SNPs on log_10_-transformed taxa abundance using a linear model. It was applied to all taxa present in ≥ 20% of individuals and SNPs with MAF ≥ 5% in the corresponding data series. The second model tested the effect of individual SNPs on the presence versus absence of the corresponding taxon using a logistic regression model. This model was applied only to taxa present in ≥ 20% and ≤ 95% of individuals and SNPs with MAF ≥ 10% (as the test statistic was inflated under the null when using this model with 5% < MAF < 10%; Suppl. Fig. 3A) (Suppl. Table 5). Both models were implemented with the GenABEL R package (Aulchenko et al., 2007) and included sex, batch and the three first genomic principal components as fixed covariates. P-values were further adjusted for residual stratification by genomic control. We obtained 1,527 signals encompassing at least three variants with *p* value ≤ 5 × 10^−8^ (the standard genome-wide significance threshold). To evaluate whether this number exceeded expectations assuming that all tests were null hypotheses, we performed two analyses. In the first we chose one of the largest (hence best powered) data series (day 240 feces in the F7 generation) and repeated all GWAS on a dataset with permuted (within litter) genotype vectors. The number of microbiota QTL (mQTL) signals detected with the real dataset was 221, while the number detected with the permuted dataset was 152, hence suggesting a true discovery rate of ~30%. In the second, we collected – for each of the 1,527 signals with p-value(s) ≤ 5 × 10^−8^ described above (corresponding each to a lead SNP x taxon x trait x cohort x model combination) – the p-values for the same SNP x taxon x model combination, yet for all other trait x other cohort combinations. Thus, we would typically collect ~5-7 p-values for each such signal. We reasoned that if the initial signals included a sufficient proportion of true positives, the lead SNPs would have similar effects in at least some of the traits in the other cohort and the collected p-values concomitantly shifted to low values. The corresponding distribution of p-values was examined by means of a QQ-plot, and was compared with the distribution obtained with an equivalent number of randomly selected series of 5-7 p-values (matched for SNP MAF and taxa abundance). This revealed a strong shift towards low p-values when compared to controls for the analyses based on abundance (rather than presence vs absence), providing additional evidence for the occurrence of real mQTL in our data (Fig. 4A). Of note, the average (F6 and F7) number of genome-wide significant mQTL was positively correlated with the average (F6 and F7) taxon’s heritability, particularly for D240 fecal samples (p = 5.2×10^−6^) (Suppl. Fig. 3B).

**Figure 4:**
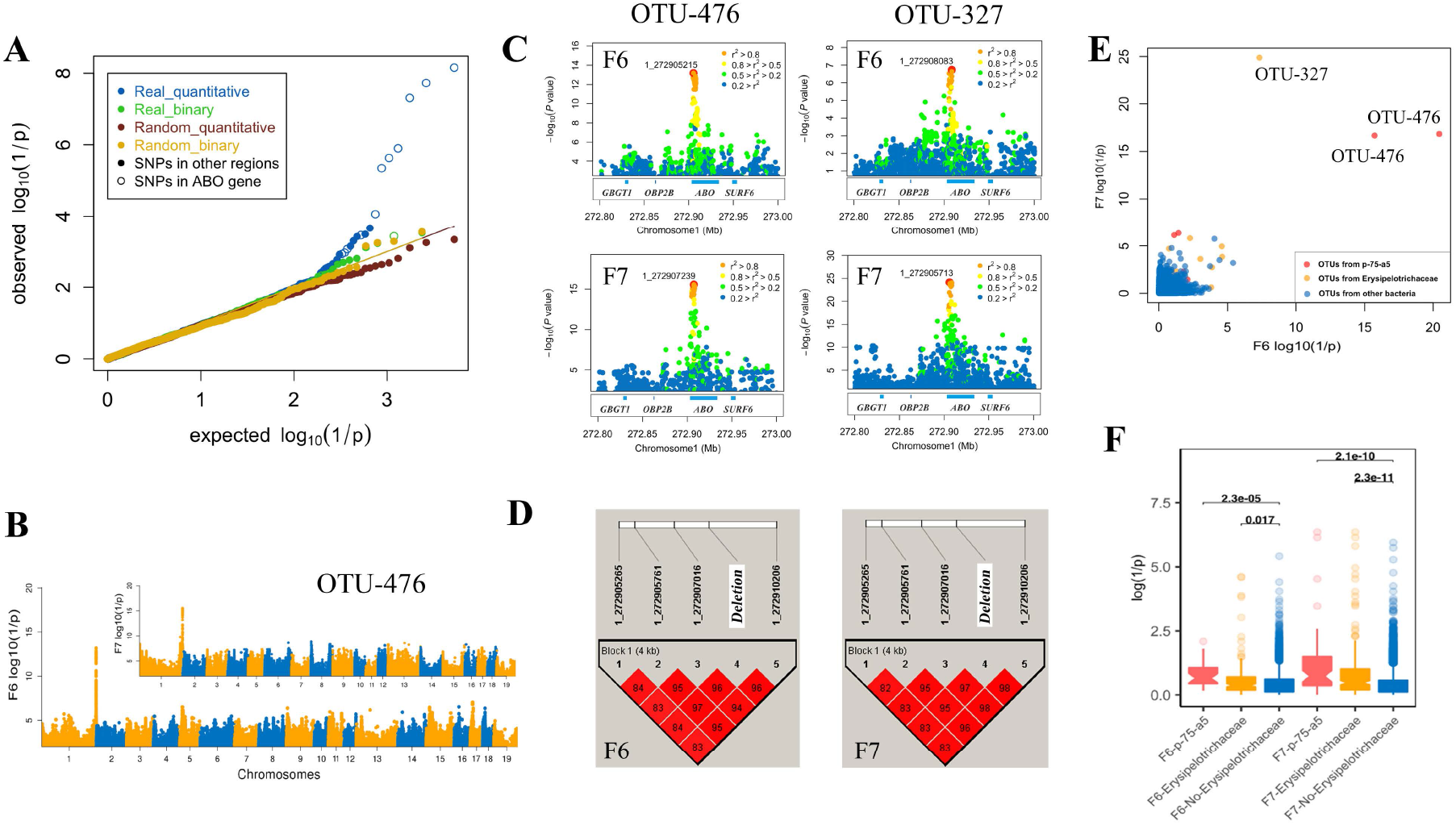
**(A)** QQ plot for 1,527 (number of signals (SNP x taxon x model x one data series in one cohort) exceeding the genome-wide log(1/p) threshold value of 7.3) sets of 5-7 ≤ p-values (same SNP x taxon x model, all data series in the other cohort) for real SNPs (Blue: quantitative model; Green: binary model), and matched sets of ≤ 5-7 p-values corresponding to randomly selected SNP x taxon combinations matched for MAF and abundance or presence/absence rate (Brown: quantitative model; Yellow: binary model). **(B)** Result of genome-wide meta-analysis in the F6 and F7 generation for OTU-476 (Manhattan plot). **(C)** Local zooms (chromosome 1: 272.8-273Mb) for OTU-476 and OTU-327 in F6 and F7. **(D)** Linkage disequilibrium (r^2^) between the four top SNPs and the 2.3Kb ABO deletion in the F6 and F7 populations (see Fig. 5). **(E)** Log(1/p) values in F6 (x-axis) and F7 (y-axis) generations for association between SNP 1_272907239 genotype and abundance of 7,748 OTUs for all studied traits and used analyses methods. OTUs that belong to p-75-a5 (respectively *Erysipelotrichaceae*) are shown in red (respectively yellow). **(F)** Comparing the distribution of association (1_272907239) p-values for p-75-a5 and *Erysipelotrichaceae* OTUs with other OTUs in F6 and F7.

To identify the corresponding mQTL yet properly accounting for the large number of realized tests before declaring experiment-wide significance and simultaneously provide confirmation in an independent cohort, we performed meta-analyses (across traits) separately in the F6 and F7 generations for the 1,527 above-mentioned signals. We designed an empirical meta-analysis approach that accounts for phenotypic correlation across traits if it exists (cfr. Methods). The discovery threshold was set at 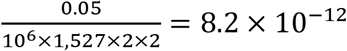 hence corrected for the size of the genome, the number of tested taxa (because genome-wide significant), the two used statistical models, and the two studied cohorts (F6 and F7). There was no need to correct for the number of traits as the meta-analysis generated one statistic for all traits. The confirmation threshold was set at 0.05/*n* where *n* was the number of signals exceeding the discovery threshold in at least one cohort. Thus, we searched for signals (defined as lead SNP x taxon x method combinations) that would exceed the discovery threshold in either the F6 or F7 cohort and the confirmation threshold in the other. We identified seven signals exceeding the discovery threshold in at least one cohort, hence setting *n*. For six of those, the confirmation threshold (i.e. 0.05/7=0.007) was also exceeded in the other cohort. All of these mapped within 3,037 bp from each other on chromosome 1 (between positions 272,904,923 and 272,907,960). They affected two individual OTUs (OTU-476 and OTU-327) as well as genus p-75-a5 to which OTU-476 is assigned (Suppl. Table 6). P-75-a5 contains 31 OTUs (other than OTU-476) that were present in ≥ 20% of samples for at least one trait. All three (OTU-476, OTU-327 and p-75-a5) are part of the *Erysipelotrichaceae* family, which contains a total of 116 OTUs subjected to GWAS. To better characterize the identified mQTL, we reran GWAS for OTU-476 and OTU-327 separately in the F6 and F7 populations (using all SNPs). The results of the corresponding association analyses are shown in Fig. 4B&C. The top SNPs on chromosome 1 (OTU-476: 1_272905215 and 1_272907239, OTU-327: 1_272908083 and 1_272905713) mapped 2,869 base pairs apart, providing a quantitative estimate of the mapping accuracy. The four SNPs were in high linkage disequilibrium with each other in both F6 and F7 populations as expected (Fig. 4D). To determine whether the corresponding mQTL might affect other taxa, we plotted the F6 and F7 association log(1/p) values for SNP 1_272907239 and the 7,748 studied OTUs. OTU-476 and OTU-327 were clearly standing out as being highly significant in both F6 and F7 (Fig. 4E). Yet the p-values for the 31 other p-75-a5 OTUs and the 83 (= 116-31-2) other *Erysipelotrichaceae* OTUs were significantly shifted towards lower p-values (Fig. 4F) with sign consistent with that for OTU-476, OTU-327 and p-75-a5 in both F6 and F7 (Suppl. Fig. 3C), suggesting that the chromosome 1 mQTL also affects other species in this family.

**Suppl. Fig. 3:**
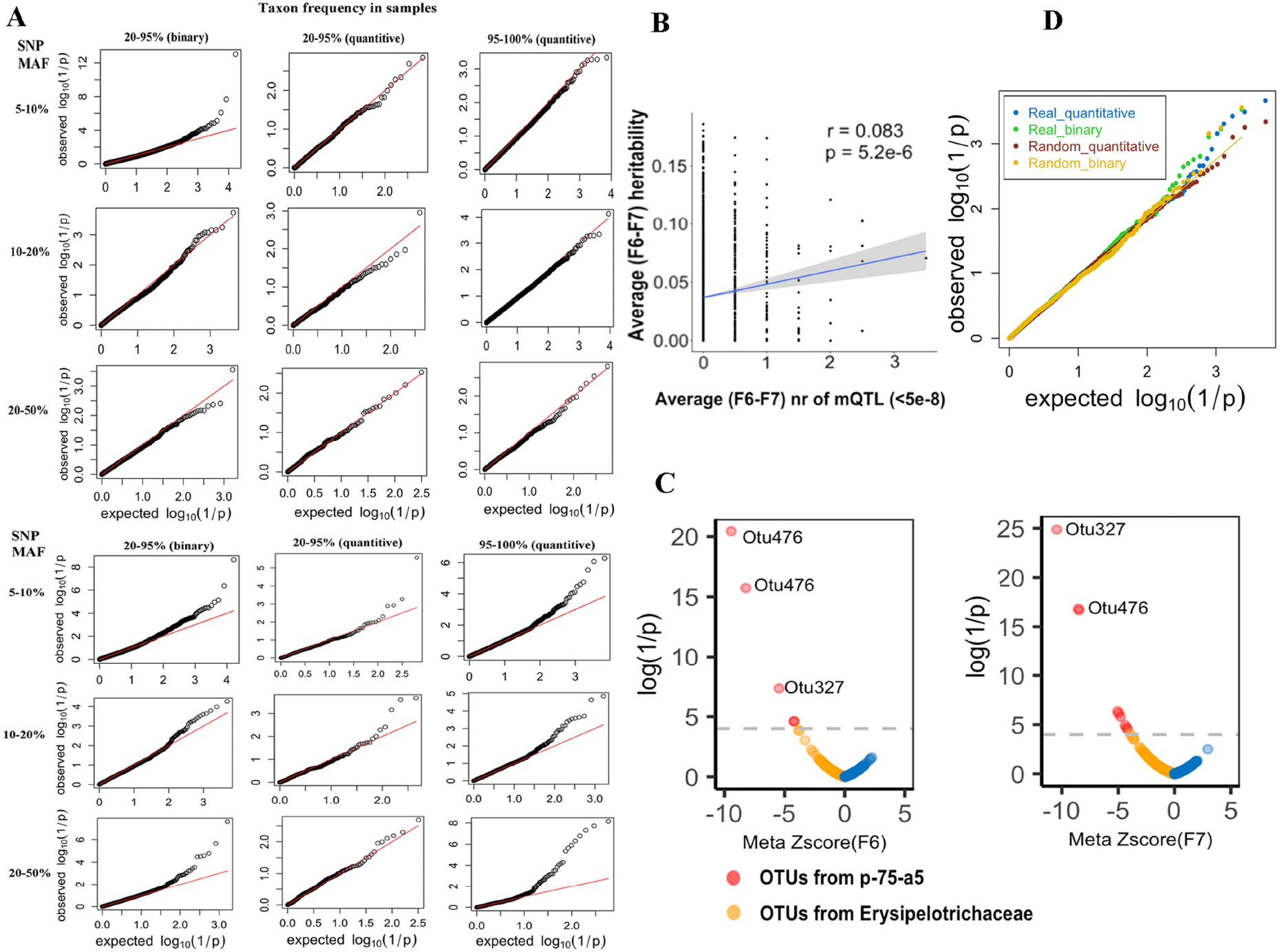
**(A) (Upper)** Distribution of log(1/p) values for 1,527 sets of 11 p-values obtained in 11 data-series for a SNP x taxon x analysis model combination that yielded a genome-wide significant signal (p < 5 x 10^−8^) in the 12^th^ data-series. **(Lower)** Distribution of log(1/p) values for 1,527 sets of 11 p-values obtained in the same data-series and with the same analysis model as in (upper) but with randomly selected SNP x taxon combinations matching the ones in (upper) for MAF and taxa abundance. **(B)** Correlation between the average (F6 and F7) taxon heritability, and the average (F6 and F7) number of genome-wide significant (*p* ≤ 5 × 10^−8^) mQTL for D240 fecal samples. **(C)** Distribution of the association log(1/p) values and corresponding signed z-scores for SNP 1_272907239 and 31 p-75-a5 OTUs (red) and 83 *Erysipelotrichaceae* OTUs, showing an enrichment of effects with same sign as for OTU-476 and OTU-327. **(D)** Same QQ plot as in Fig. 4A after removal of all SNPs in the chromosome 1: 272.8-273.1Mb interval.

**Supplemental Table 5:** Number of SNPs and taxa used for mQTL analyses in the different data series

| Generation | Data series | Sample type | Sample size | Nr SNPs<br>(MAF ≥ 5%) | Quantitative model | | Binary model | |
| --- | --- | --- | --- | --- | --- | --- | --- | --- |
| | | | | | Nr taxa<br>(20% $< F < 95\%$) | Nr taxa<br>($F \geq 95\%$) | Nr SNPs<br>(MAF ≥ 10%) | Nr taxa<br>($F > 20\%$) |
| F6 | D25 | Feces | 81 | 25,725,345 | 1,372 | 132 | 20,477,556 | 1,372 |
|  | D120 | Feces | 475 | 26,572,513 | 2,519 | 372 | 20,814,322 | 2,519 |
|  | D240 | Feces | 633 | 26,634,424 | 3,246 | 375 | 20,761,630 | 3,246 |
|  | IC | Ileal content | 288 | 26,660,960 | 869 | 68 | 20,733,869 | 869 |
|  | CC | Cecum content | 292 | 25,513,317 | 1,983 | 275 | 20,698,139 | 1,983 |
| F7 | D25 | Feces | 232 | 26,517,529 | 1,731 | 95 | 20,730,691 | 1,731 |
|  | D120 | Feces | 405 | 26,540,167 | 4,055 | 290 | 20,694,289 | 4,055 |
|  | D240 | Feces | 582 | 26,562,071 | 4,124 | 349 | 20,645,179 | 4,124 |
|  | IC | Ileal content | 408 | 26,623,600 | 499 | 55 | 20,677,324 | 499 |
|  | IM | Ileal mucosa | 76 | 25,756,486 | 1,606 | 256 | 20,356,739 | 1,606 |
|  | CC | Cecal content | 637 | 26,632,837 | 3,762 | 205 | 20,652,282 | 3,762 |
|  | CM | Cecal mucosa | 483 | 26,591,809 | 1,664 | 225 | 20,676,604 | 1,664 |

**Supplemental Table 6:** Signals exceeding the experiment-wise significance threshold in at least one cohort.

| Taxon | SNP | meta-p-value |  |
| --- | --- | --- | --- |
|  |  | F6 | F7 |
| Otu327(f_Erysipelotrichaceae) | 1_272907960 | 2.55E-06 | 5.24E-24 |
| g_p-75-a5 | 1_272907573 | 5.84E-04 | 3.81E-17 |
| g_p-75-a5 | 1_272907165 | 3.05E-12 | 2.76E-16 |
| Otu16389(k_Bacteria) | 6_169188720 | 6.19E-01 | 1.78E-15 |
| Otu476(g_p-75-a5) | 1_272907573 | 1.47E-04 | 4.57E-15 |
| Otu476(g_p-75-a5) | 1_272907165 | 1.12E-13 | 1.38E-14 |
| Otu476(g_p-75-a5) | 1_272904923 | 1.54E-14 | 8.76E-14 |

(showing top signals only)

### The chromosome 1 mQTL is caused by a 2.3-Kb deletion in the ABO acetyl-galactosaminyl-transferase gene that is under balancing selection

All lead SNPs of the meta-analyses conducted in the F6 and F7 generation map to the 3’ end of the porcine acetyl-galactosaminyl transferase gene that is orthologous to the gene underlying the ABO blood group in human (Fig. 4B). This is a strong candidate gene known to modulate interactions with several pathogens (Cooling et al., 2015), although not directly known to affect intestinal microbiota composition in healthy humans (Davenport et al., 2016). Four non-synonymous ABO SNPs (*R37G, A48P, S60R, G66C*) segregated in the F6 and F7 generation but none of these were in high LD with any of the lead SNPs (*r*^2^ ≤ 0.18). A 2.3 Kb deletion encompassing the last exon (eight) of the acetyl-galactosaminyl transferase gene has been previously reported in the pig. It causes a null allele equivalent to human “O”, while the wild-type allele corresponds to the human “A” allele with alpha 1-3-N-acetyl-galactosaminyl-transferase activity (Choi et al., 2018). The pig *Sus scrofa* 11.1 reference genome corresponds to the “O” allele. We de novo assembled an “A” allele using PacBio whole genome sequence data from one of our Bamaxiang animals. We confirmed the boundaries of the 2.3 Kb deletion and showed that it results from an intra-chromosomal recombination between SINE elements (Fig. 5A and Suppl. Fig. 4A). The four top SNPs in Fig. 4D mapped within 2.1 Kb from the 2.3Kb deletion. We developed a PCR test and genotyped all F0, F6 and F7 animals for the “O” deletion. Three of the four top SNPs were in perfect LD with the deletion in the F0 generation and near-perfect (*r*^2^ ≥ 0.94) in the F6 and F7 generations (Fig. 4D). None of the four non-synonymous ABO variants had an independent effect on OTU-476 or 327 abundance. We mapped cecal RNA-Seq data from AA, AO and OO individuals on the Bamaxiang A reference allele. Transcripts from the AA individuals showed the expected splicing pattern yielding an ~1.25 Kb mRNA coding for 364 amino-acid of which 230 (63%) by exon 8. Transcripts from OO individuals were characterized by the use of an alternative 70 bp eighth and 6.9 Kb ninth exon flanked by canonical splice sites (Fig. 5A). The corresponding 7.4 Kb mRNA substitutes the 230 amino-acids encoded by the wild-type eight exon with a shortened lysine-serine-isoleucine carboxyterminal tail. The encoded truncated protein misses seven of the eight substrate binding sites and seven of the eight active sites reported by Wang et al. (2015). The proportion of reads mapping to the seventh intron was higher for the O than for the A allele pointing towards less efficient splicing of the alternative intron. We used three synonymous variants in LD with the 2.3 Kb deletion (mapping respectively in exons 4, 6 and 7) to measure allelic imbalance from RNA-Seq data of AO individuals. O transcripts accounted for 26% of acetyl-galactosaminyl transferase transcripts in AO individuals, possibly reflecting non-sense mediated RNA decay due to a stop codon in the penultimate exon (Fig. 5 and Suppl. Fig. 4B). This ~3-fold reduction in abundance of O versus A transcripts was confirmed by expression QTL (eQTL) analysis performed using RNA-Seq data from 300 F7 cecum tissues samples (p =1.9×10^−43^) (Suppl. Fig. 4C). Taken together, our results indicate that the 2.3 Kb “O” deletion is a null allele and the most likely mQTL causative mutation.

**Figure 5:**
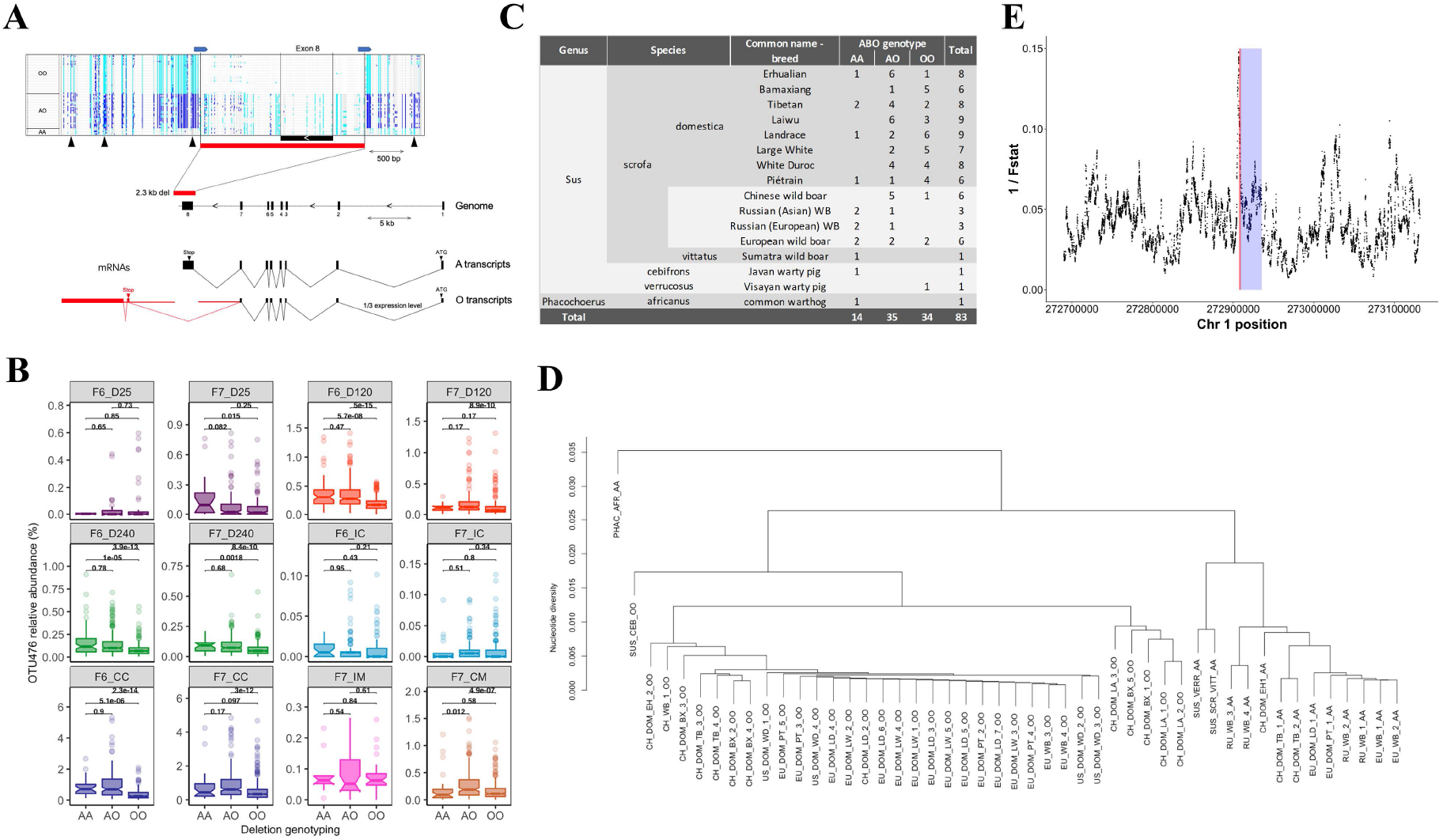
**(A) Structure of the porcine ABO acetyl-galactosaminyl transferase gene with position of the 2.3 Kb deletion (red rectangle)**. Screen capture of Integrated Genome Viewer (IGV) view of the genotypes of the 61 F0 animals (sorted by OO, AO and AA genotype) for 145 variants in a ~5 Kb interval spanning the 2.3 Kb deletion. Sequence reads were mapped to the Bamaxiang A allele as reference. Light blue: homozygous for alternate allele; dark blue: heterozygous alternate/reference; gray: homozygous for reference allele. The horizontal blue arrows mark the position of SINE sequences that may have mediated the intra-chromosomal recombination event that has created the 2.3 Kb deletion. The vertical black arrow mark the position of the top variants reported in Fig. 4C. Effect of the 2.3Kb deletion on the structure and abundance of acetyl-galactosaminyl transferase transcripts: (i) creation of alternate exon 8 and 9, and (ii) reduction of transcript levels to ~ 1/3th of normal levels. **(B) Effect of acetyl-galactosaminyl transferase genotype (AA, AO or OO) on abundance of OTU-476 in the twelve data series** showing that (i) the effect of the A allele is dominant over that of the O allele, and (ii) the mQTL effect is detected in cecum (content and mucosa) and in day 120 and 240 feces. **(C) The AO acetyl-galactosaminyl transferase polymorphism is a trans-species polymorphism in Suidae**. Distribution of the AO genotype in domestic *S. scrofa* (domestic pigs), wild *S. scrofa* (wild boars), *S. verrucosus* (Visayan warty pig), *S. cebifrons* (Javan warty pig), and *Phacochoerus Africanus* (common warthog). **(D)** UPGMA dendrogram based on sequence similarity between the chromosomes of 14 homozygous AA and 34 OO animals in a 5-Kb window centered around the 2.3Kb deletion (variants inside the deletion were ignored). PHAC_AFR: common warthog, SUS_VERR: Visayan warty pig, SUS_CEB: Javan warty pig, SUS_SCR_VII: Sumatran wild boar, CH/RU/EU_WB: Chinese/Russian/European wild boars, CH/EU/AM_DOM: Chinese/European/American domestic pigs. Breed acronyms are as in Fig. 1. **(E) Peak of reduced population differentiation between eight domestic breeds coinciding with the 2.3 Kb deletion (red) in the porcine acetyl-galactosaminyl transferase gene (blue)**. X-axis: position on porcine chromosome 1. Y-axis: 1/(mean F statistic) for all variants in a 2Kb sliding window. F statistic computed as the ratio of the “between-breed mean squares” and the “within-breed mean squares” for the dosage of O allele.

We closely examined the effect of AO genotype on the abundance of the affected OTUs (OTU-476, OTU-327 and p-75-a5) in the 12 data series. This clearly showed (i) that the effect of the A allele is dominant over that of the O allele, and (ii) that the effect manifests in D120 and D240 feces, cecal content as well as mucosa, but not in D25 feces, ileal content and mucosa (Fig. 5B and Suppl. Fig. 4D). In these samples (D120, D240, CC, CM), AO genotype explained on average 7.9%, 3.2% and 6.6% of the variance in abundance for OTU-476, OTU-327 and genus p-75-a5 (Suppl. Fig. 4E). Of note, the abundance of OTU-476 and OTU-327 was shown to be highest in cecal content where they account on average for respectively ~0.92% and ~0.02% of reads in AA/AO animals, and for 0.47% and 0.003% of reads in OO animals (Fig. 5 and Suppl. Fig. 4F).

The ABO locus is known in humans to be under strong balancing selection that has perpetuated identical-by-descent alleles segregating in present humans, gibbons and Old-World monkeys for tens of millions of years (Ségurel et al., 2012). To verify whether a similar situation might occur in pigs, we analyzed the sequences of the 61 F0 animals (*Sus scrofa domestica*), 15 wild boars (9 Asian, 7 European)(*Sus scrofa*), one Indonesian wild boar from Sumatra (*Sus scrofa vittatus*), one Visayan warty pig from the Philippines (*Sus cebifrons*), one Javan warty pig from Indonesia (*Sus verrucosus*), and one common warthog from Africa (*Phacochoerus africanus*) in a 50 Kb window spanning the ABO gene. Asian and European wild boar (and derived domestic breeds) are thought to have diverged from a common *Sus scrofa* ancestor ~1 million years ago (MYA), *Sus scrofa* and *Sus scrofa vittatus* ~1.5 MYA, *Sus scrofa* and *Sus cebifrons/verrucosus* ~3.5 MYA, and *Sus scrofa* and *Phacochoerus africanus* ~10 MYA (Groenen, 2016). The same (identical breakpoints) 2.3Kb deletion was shown to segregate in all eight F0 breeds, in all Asian and European/American wild-boar populations, and – remarkably – in *Sus cebifrons* (Fig. 5C). Consistent with the hypothesis of a trans-species polymorphism (rather than hybridization), the O allele of *Sus cebifrons* was shown to lie outside of the cluster of *Sus scrofa* O alleles (Fig. 5D). Further supporting the hypothesis of balancing selection, the ABO gene was characterized by a marked drop in population differentiation between domestic pig breeds maximizing exactly at the position of the 2.3 Kb deletion (Fig. 5E).

**Suppl. Fig. 4:**
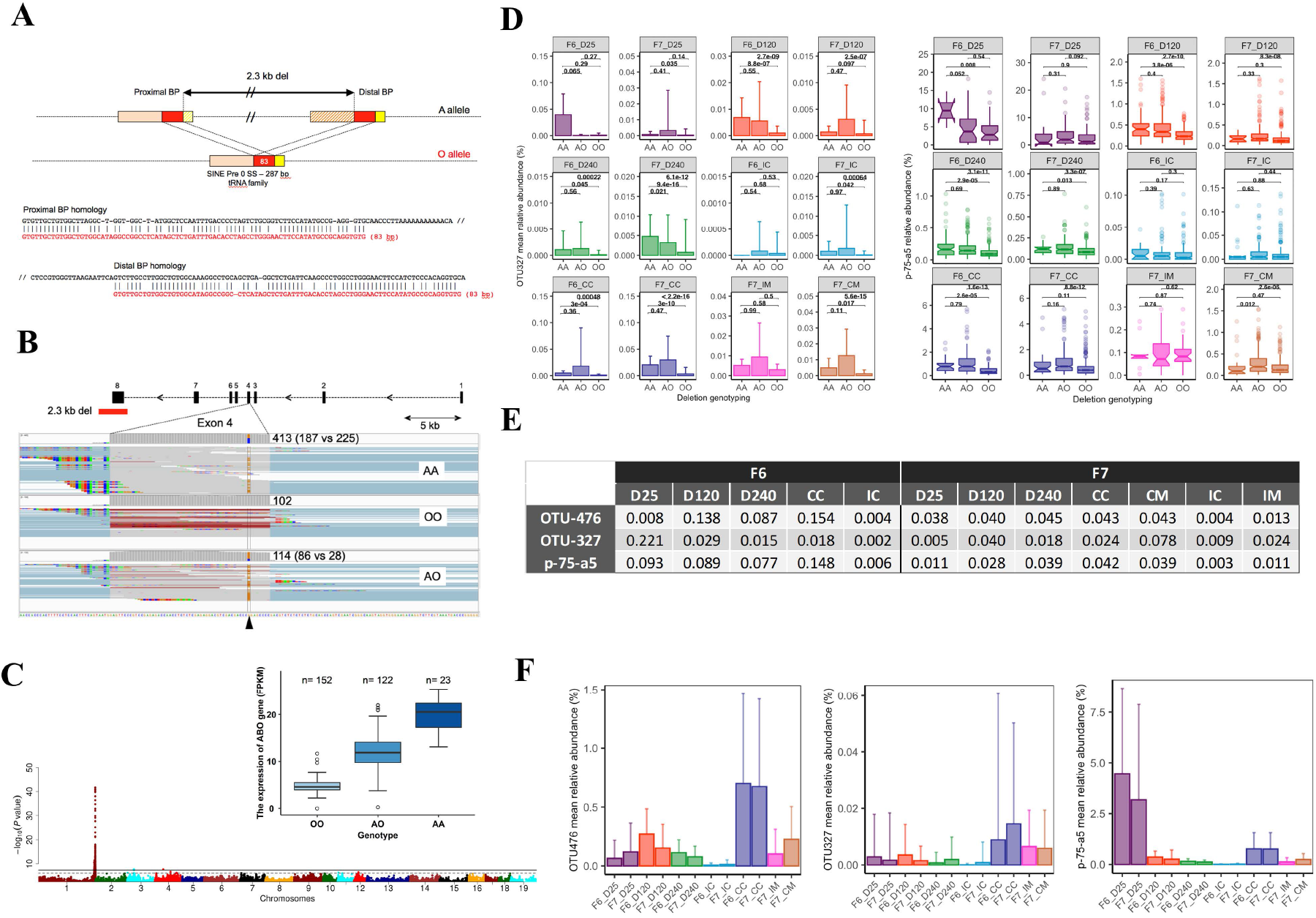
**(A)** Breakpoints of the 2.3 kb deletion showing the role of a duplicated SINE sequence in mediating an intra-chromosomal recombination. **(B)** Illustrative example of allelic balance for the *cG146C* SNP in an AA homozygote and of allelic imbalance for the same SNP in an AO heterozygote. **(C)** eQTL analysis for the ABO gene maximizing at the exact position of the 2.3Kb deletion (p = 1.9×10^−43^) and showing the additive effect of the A allele increasing transcript levels ~3-fold (inset; FPKM: Fragments Per Kilobase of transcript per Million mapped reads). **(D)** Effect of acetyl-galactosaminyl transferase genotype (AA, AO or OO) on abundance of OTU-327 and p-75-a5 in the twelve data series. **(E)** Fraction of the variance in abundance of the corresponding OTU/genus explained by AO genotype. **(F)** Abundance of OTU-476, OTU-327 and p-75-a5 in the twelve data series.

### The chromosome 1 mQTL affects bacterial species with complete N-acetyl-D-galactosamine (GalNAc) import and catabolic pathway

In human, the ABO acetyl-galactosaminyl transferase gene is broadly expressed yet particularly strongly in the small and large intestine (Suppl. Fig. 5A). We characterized the expression profile of the porcine ABO gene in a panel of 15 tissues in an adult animal (Bamaxiang sow) and a fetus (Duroc male) by RNA-seq. A very similar expression profile was observed in the pig with strong expression in the gastrointestinal tract, particularly in the adult (Suppl. Fig. 5B). The acetyl-galactosaminyl-transferase encoded by the A allele adds GalNAc (*α*1-3 linkage) to a variety of glycan substrates sharing a Fuc*α*1-2Gal/*β*1-4GlcNAc or Fu*α*1-2Gal/*β*1-3GlcNAc (H antigen) extremity (Cooling, 2015). In the gut, these include the heavily glycosylated secreted and transmembrane mucins constituting the cecal mucus. Mucin glycans are used as carbon source by the intestinal microbiota, especially under low-fiber diet (Ravcheev & Thiele, 2017; Zuniga et al., 2018). We reasoned that the observed mQTL might act by altering the intestinal concentration of GalNAc, the A allele thereby favoring the growth of bacterial species effective at utilizing this sugar.

To test this hypothesis, we first measured the concentration of GalNAc in cecal content by LC-MS/MS in 17 AA animals and 17 OO animals of the F7 generation. GalNAc concentrations were indeed ~1.8-fold higher in AA than in OO pigs (p = 5.6 x 10^−4^) (Fig. 6A).

**Figure 6:**
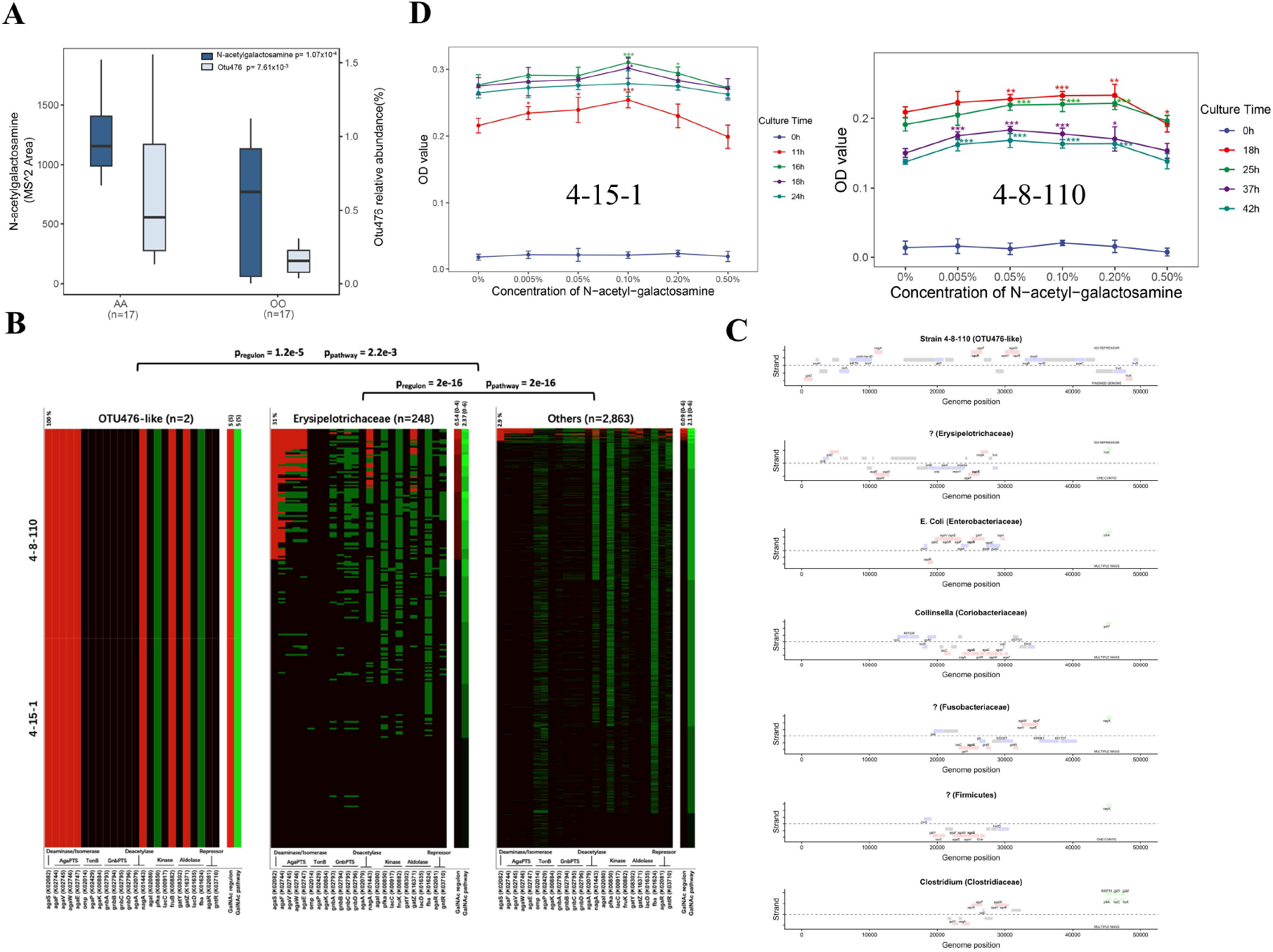
**(A)** Concentrations of GalNAc measured by LC-MS/MS in cecal content of 17 AA and 17 OO day 240 pigs. Abundances of OTU-476 determined by 16S RNA gene sequencing are shown for the same samples. **(B)** Presence anywhere in the genome (green), presence in close proximity to *agaS* (red), or absence (black) of the orthologues of 24 genes implicated in the GalNAc TR/CP pathway in the genome of (i) two OTU-476 like strains (4-15-1 and 4-8-110), (ii) 248 MAGs assigned to the *Erysipelotrichaceae* family, and (iii) 2,863 MAGs assigned to other bacterial families. The two lanes on the right of the three panels correspond to the Regulon (red) and Pathway (green) score respectively. Both scores range from 0 (black) to 6 (bright red or green). Means (range) for the corresponding dataset are given on top. **(C)** Maps of GalNAc “operons” in one of the two OTU476-like strains and six MAGs assigned respectively to an *Erysipelotrichaceae*, *E. coli* (an *Enterobacteriaceae*), a *Collinsella* (a *Coriobacteriaceae*), a *Fusobacteriaceae*, a *Firmicutes* and a *Clostridium*. Identified Open Reading Frames (ORFs) are represented as colored boxes. Genes implicated in GalNAc import and catabolism are in red if they are part of the cluster and in green if located elsewhere in the genome. Genes with a known function unrelated to GalNAc are in blue. ORFs with uncharacterized gene product in gray. Gene acronyms are given next to the corresponding boxes. ORFs transcribed from the top (respectively bottom) strand are above (below) the dotted line. The source of information used to confirm the map order is given (finished genome, multiple MAGs, single contig). **(D)** Growth curves (0 to 42 hours) of OTU476-like strains 4-15-1 and 4-8-110 in the presence of growing concentrations of GalNAc. *: p< 0.05, **: p<0.01, ***: p<0.001. The p-values correspond to the difference between the growth rate with and without addition of GalNAc at the same time.

To gain insights in the relative capacity of porcine intestinal bacteria to utilize GalNAc, we then (i) isolated two bacterial strains (4-8-110 and 4-15-1) with V3-V4 sequence similarity of 100% and 99.8% with OTU-476 from porcine feces and sequenced their genome on a ONT PromethION platform (Oxford Nanopore Technology, UK), and (ii) built 3,111 metagenomic assembled genomes (MAGs) from shotgun sequence data obtained from 92 samples including feces, content of three intestinal locations (jejunum, ileum, cecum), and eight populations (26 F6, 12 Duroc, 12 Large White, 12 Tibetan, 6 Laiwu, 6 Licha, 6 Berkshire x Lisha F1s, and 12 Chinese wild boars). Of the 3,111 MAGs, 248 were assigned to the family *Erysipelotrichaceae* using PhyloPhlAn (Segata et al., 2013). To be used as carbon source by intestinal bacteria, GalNAc needs (i) to be released from the glycan structures by secreted glycosyl hydrolases (GH), (ii) to be imported across the bacterial membranes by dedicated transport systems (TR), and (iii) to be converted into intermediates of central metabolism by a specific catabolic pathway (CP). While some bacteria may have both GH and TR/CP for specific monosaccharides, other may only have the GH (“donors”) or the TR/CP (“acceptors”) (Ravcheev & Thiele, 2017). We compiled a list of 24 genes (with corresponding KEGG ontology number) implicated in GalNAc utilization (TR/CP) from the literature (Brinkkötter et al., 2000; Rodionov et al., 2010; Leyn et al., 2012; Hu et al., 2012; Biddart et al., 2014; Zhang et al., 2015; Ravcheev & Thiele, 2017, Zuniga et al., 2018). These encode (i) 11 components of one of three GalNAc transporter systems (AgaPTS: *agaE, agaF, agaV, agaW*; TonB dependent transporter: *omp, agaP, agaK*; GnbPTS: *gnbA, gnbB, gnbC, gnbD*), (ii) two GalNAc-6P deacetylases (*agaA, nagA*), (iii) two galactosamine-6P (GalN-6P) isomerase and/or deaminases (*agaI, agaS*), (iv) three tagatose-6P kinases (*pfka, lacC, fruK*), (v) four tagatose-1,6-PP aldolases or aldolase subunits (*gatY-kbaY, gatZ-kbaZ, lacD, fba*), and (vi) two regulon repressors (*agaR, gntR*), for a total of six essential pathway constituents (Suppl. Table 7). Genes involved in the utilization of specific sugars (including GalNAc) tend to cluster and form operons of potentially coregulated genes (regulons) that support all or most of the essential TR/CP steps. The steps that are not encoded by the operon may be complemented in trans by genes encoding enzymes that are often less substrate-specific (Lawrence, 1999; Koonin, 2009). We used GhostKOALA (Kanehisa et al., 2016) to search for orthologues of the 24 genes in the two OTU476-like genomes and 3,111 MAGs. We generated two scores to evaluate the capacity of bacterial species to utilize GalNAc. The first (pathway score) counted the number of essential steps in GalNAc utilization (out of six) that could be accomplished by the set of orthologues detected in the genome (cfr. Methods), irrespective of their map position. The second (regulon score) counted the number of essential GalNAc utilization steps that could be fulfilled by orthologues that were clustered in the genome, i.e. forming a potential operon. Following Ravcheev and Tiele (2017), we used *agaS* as anchor gene to establish the regulon score, i.e. we counted how many essential steps in GalNAc utilization (out of six) were covered by genes located in the vicinity of *agaS*.

The first striking observation was that at least one orthologue of *agaS* was found in the two (=100%) OTU476-like strains (4-15-1 and 4-8-110), in 31% of *Erysipelotrichaceae* MAGs (n=248), yet in only 3.0% of other MAGs (n=2,863). The second, was that both scores (pathway and regulon score) were very significantly higher for *Erysipelotrichaceae* than for other MAGs (p_pathway_=2.0e-16 and p_regulon_=2.0e-16), and for the two OTU476-like strains than for *Erysipelotrichaceae* and non-*Erysipelotrichaceae* MAGs combined (p_pathway_=2.2e-3 and p_regulon_=1.2e-5) (Fig. 6B). These comparisons accounted for variation in the MAGs’ completion score, number of contigs and predicted size of the corresponding genomes (see Methods). Examination of the genome of the two OTU476-like strains revealed clustering of eight GalNAc genes including orthologues of the four components of the AgaPTS transporter system (*agaE, agaF, agaV, agaW*), of *nagA* deacetylase, of *agaS* deaminase/isomerase, of *fruK* kinase, and of the *gatZ-kbaZ* aldolase subunit. This amounted to a score of five for both pathway and regulon score, corresponding (after accounting for completion, contig number and genome size) to the top 4.7% and 0.35% of 3,113 pathway and regulon scores, respectively. The organization of the GalNAc gene cluster was identical in both strains (4-15-1 and 4-8-110), covering ~50Kb. Intriguingly, closer examination of the corresponding region also revealed an orthologue of the *nagB* GlcNAc deaminase/isomerase, and of the *fruR2* member of the DeoR family of transcriptional regulators, which are paralogues of *agaS* and *agaR*, respectively (Fig. 6C).

We further showed that adding GalNAc in the culture medium indeed enhances the growth of the two isolated OTU-476 like strains, indicating that these can indeed utilize GalNAc as carbon source (Fig. 6D).

Taken together, these findings provide strong support for our hypothesis, i.e. that the mQTL acts by increasing cecal GalNAc concentration hence favoring the growth of bacterial species effective at utilizing GalNAc.

**Suppl. Fig. 5:**
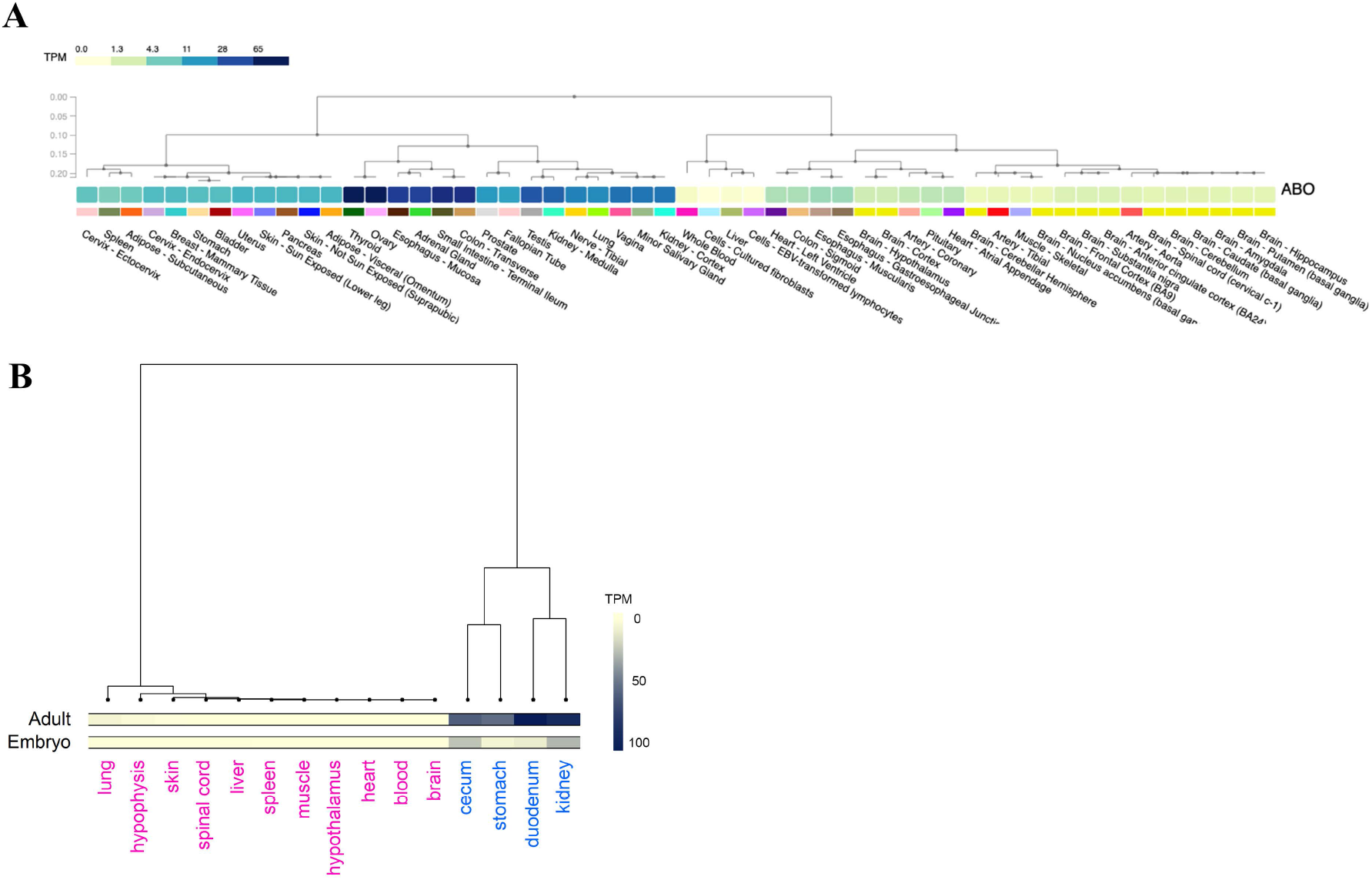
**(A)** Expression profile of the ABO gene in human tissues (from GTEx Portal: https://gtexportal.org/home/). **(B)** Expression profile in a panel of adult and embryonic porcine tissues (own RNA-Seq data).

**Supplemental Table 7:**
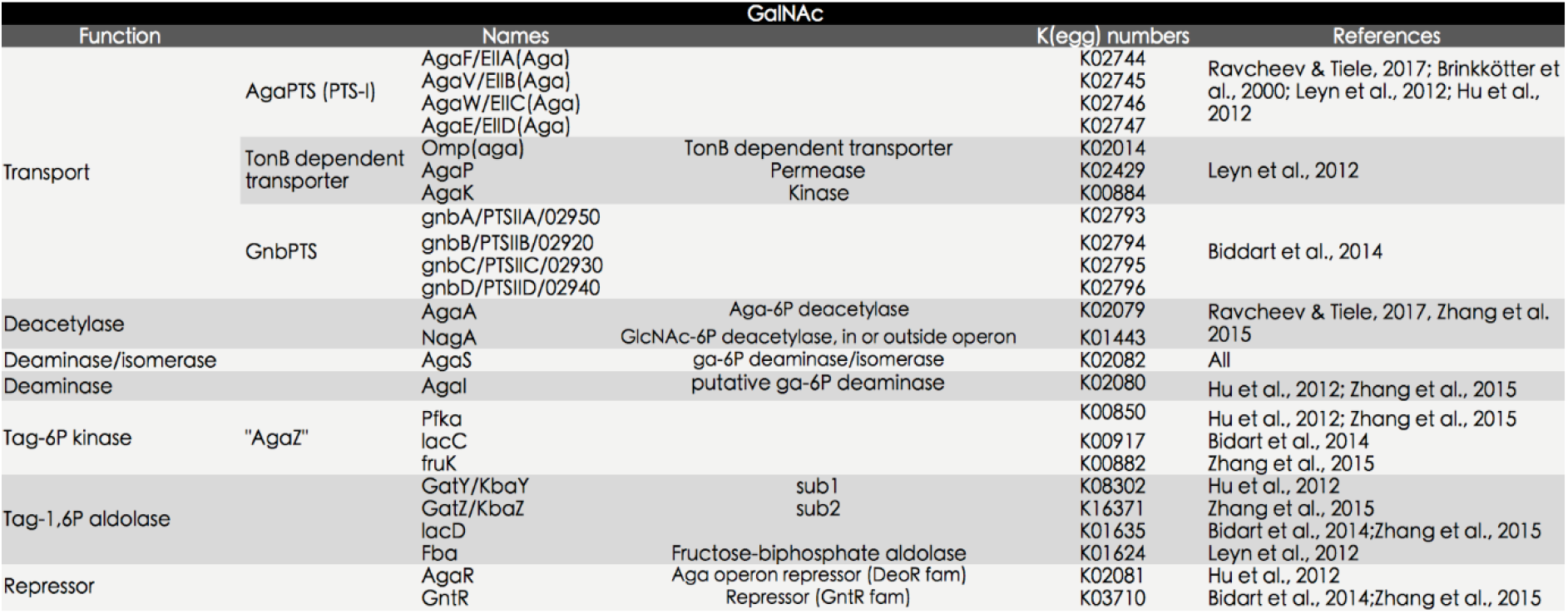
Bacterial genes implicated in GaLNac import and catabolism

### No effect of ABO genotype on intestinal microbiota composition in human

The effect of ABO genotype on intestinal microbiota composition in humans remains somewhat controversial. Despite suggestive evidence in a small (n=71) cohort of separate microbiota-based clustering of AB and B vs A and O individuals (secretors only) (Makivuokko et al., 2012), a subsequent study conducted in a larger cohort (n=1,503) could not detect experiment-wide significant effects of either secretor or ABO genotype on gut microbiota composition (Davenport et al., 2016). Intriguingly, the latter study nevertheless reported a possible effect of ABO genotype on a rare OTU (592616) assigned to *Erysipelotrichaceae*.

We took advantage of an available intestinal 16S rRNA dataset of ~300 healthy individuals of European descent to re-examine this question in light of the results obtained in the pig (Momozawa et al., 2018). All individuals were genotyped with the OmniExpress SNP array (Illumina) and imputed to whole genome. ABO genotype was inferred from the genotypes at three coding variants (rs8176719, rs7853989, rs8176747) following Cooling (2015). The frequency of the different genotypes in the cohort were: 0.37 (OO), 0.37 (AO), 0.11 (BO), 0.08 (AA) and 0.06 (AB). Twenty-one percent of the individuals in the cohort were non-secretors (homozygous for the *W143X* mutation in the *FUT2* gene) (Kelly et al., 1995). For each individual we obtained V1-V2, V3-V4 and V5-V6 16S rRNA sequences from intestinal biopsies (cfr. Methods). 16S rRNA sequences were clustered in OTUs using DNACLUST (Ghodsi et al., 2011) with a 97% similarity threshold. Forty-three (V1-V2), 20 (V3-V4) and nine (V5-V6) OTUs with abundance >0.001% across locations and amplicons were assigned to *Erysipelotrichaceae* using the Silva database (Quast et al., 2013). The effect of ABO blood group on OTU abundance was tested using a linear model including blood group (AA, AO, AB versus rest, BB, BO, BB versus rest, OO versus rest), secretor status, sex, age, smoking status and BMI (cfr. Methods). There was no convincing evidence for an effect of ABO blood group on the abundance of *Erysipelotrichaceae* OTU (Suppl. Fig. 6). The SILVA database does not include genus p-75-a5. We directly mapped the 16S rRNA reads to the Greengenes database (DeSantis et al., 2006). Putative p-75-a5 reads were detected in five individuals only (four OO and one AO) in which they accounted for 0.003%-0.25% of the reads.

**Suppl. Fig. 6:**
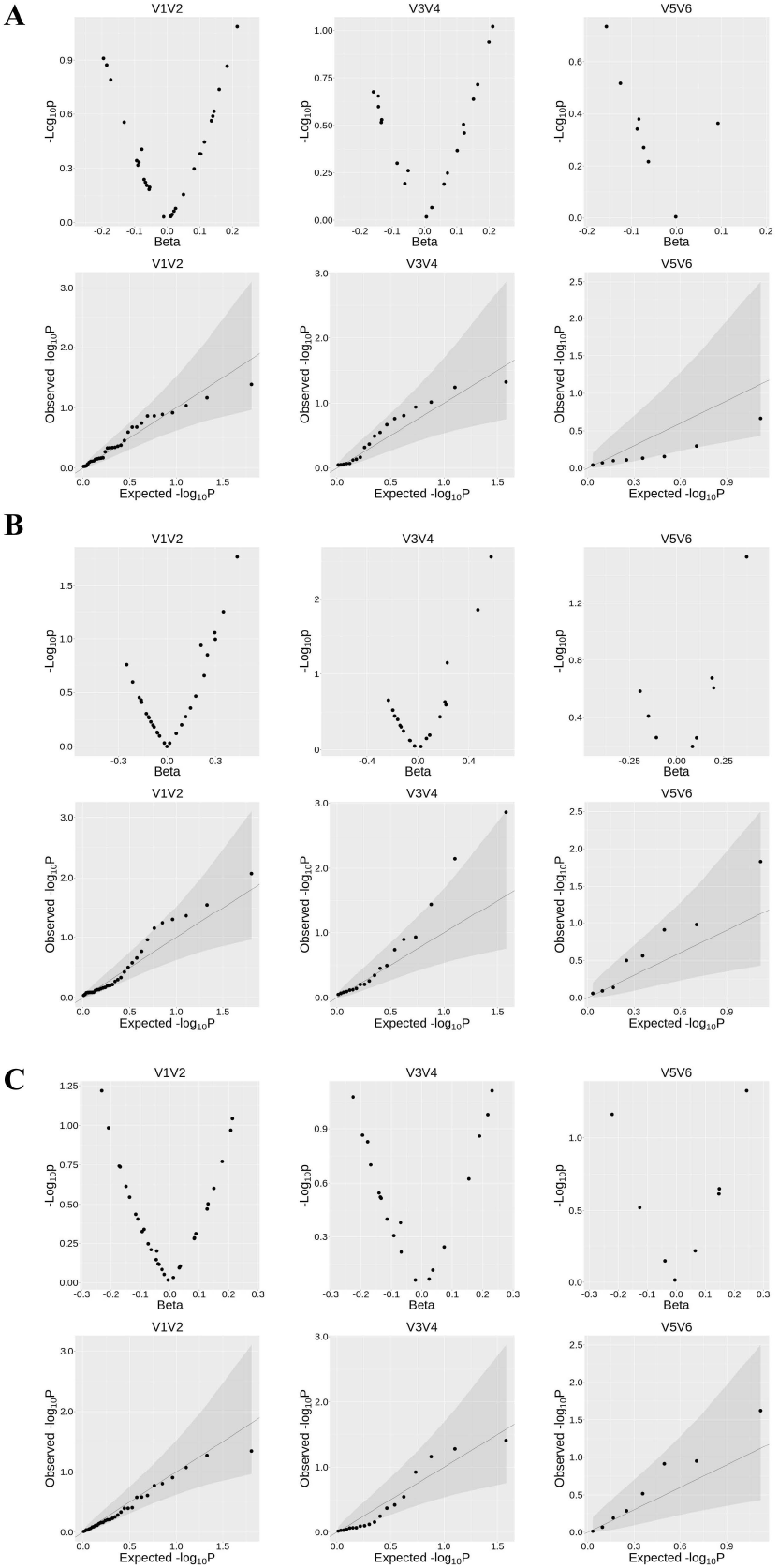
Volcano and QQ plots for 43 (V1-V2), 20 (V3-V4) and 9 (V5-V6) OTUs classified as *Erysipelotrichaceae* for the contrasts **(A)** [AA, AO and AB] versus [BB, BO and OO], **(B)** [BB, BO and AB] versus [AA, AO and OO], and **(C)** [OO] versus [all others].

## Discussion

We herein report the use of a genetically heterogeneous population to study the impact of host genetics on the composition of the intestinal microbiota of the pig. More than 30 million variants with MAF ≥ 3% segregate in this population, i.e. more than one variant every 100 base pairs. This is slightly lower than the 40 million high quality variants segregating in the mouse collaborative cross (Srivastava et al., 2017). The average nucleotide diversity (*π*, i.e. the proportion of sites that differ between two chromosomes sampled at random in the population(s)) within the four Chinese founder breeds was ~2.5×10^−3^ and within the four European founder breeds ~2.0×10^−3^. By comparison, *π*-values in African and Asian/European human populations are ~ 9×10^−4^ and ~ 8×10^−4^, respectively (Yu et al., 2001; The 1,000 Genomes Project Consortium, 2010). Thus, against intuition (as domestication is often assumed to have severely reduced effective population size) the within population diversity is > 2-fold higher in domestic pigs than in human populations, as previously reported (Frantz et al., 2015; Charlier et al., 2016; Georges et al., 2019). Nucleotide diversities between Chinese and between European founder breeds were ~3.6×10^−3^ and ~2.5×10^−3^, i.e. 1.44-fold and 1.25-fold higher than the respective within breed *π*-values. These *π*-values are of the same order of magnitude as the sequence divergence between *Homo sapiens* and Neanderthals/Denosivans (~3×10^−3^, Sankararaman et al., 2014). By comparison, *π*-values between Africans, Asians and Europeans are typically ≤ ~ 1×10^−3^ (Yu et al., 2001). The nucleotide diversity between Chinese and European breeds averaged ~4.3×10^−3^. This *π*-value is similar to the divergence between *M. domesticus* and *M. castaneus* (Geraldes et al., 2008), and close to halve the ~1% difference between chimpanzee and human (Patterson et al., 2006). Note that Chinese and European pig breeds are derived from Chinese and European wild boars, respectively, which are thought to have diverged ~1 million years ago (Groenen, 2016), while *M. domesticus* and *M. castaneus* are thought to have diverged ≤ 500,000 years ago (Geraldes et al., 2008). The genomic contribution of the eight founder breeds in the F6 and F7 generation is remarkably uniform and close to expectations (i.e. 12.5%) both at genome-wide and chromosome-wide level (Fig. 1C), suggesting comparable levels of genetic diversity across the entire genome. This does not preclude that more granular examination may reveal local departures from expectations, or under-representation of incompatible allelic combinations at non-syntenic loci. Such analyses are beyond the scope of this study.

Average microbiota composition of the 12 data-series indicates a remarkable consistency for the same traits across the F6 and F7 generation, yet marked compositional differences between traits (Fig. 2B). Even at family-level, some taxa are found to be nearly trait-specific (Suppl. Fig. 1C). For instance, the proteobacteria *Enterobacteriaceae*, *Pseudomonadaceae, Pasteurellaceae*, the firmicutes *Clostridiaceae*, *Peptostreptococcaceae*, *Bacillaceae*, *Leuconostocaceae*, and the actinobacteria *Microbacteriaceae* are at least ten times more abundant in ileal than in any other sample type. Amongst those, *Leuconostocaceae* are nearly digesta-specific, while *Pseudomonadaceae* are nearly mucosa-specific. The Bacteroidetes *Odoribacteraceae* and *Rikenellaceae* were found to be at least ten times more abundant in day 25 feces than in any other sample type. The firmicutes *Christensenellaceae* were nearly ten times more abundant in feces (irrespective of age) than in any other sample type. This confirms that limiting the analysis of the intestinal microbiota to adult fecal samples can only provide a very partial view of its complexity and the factors that determine it (Donaldson et al., 2016).

To evaluate the importance of host genetics in determining gut microbiota composition we first examined the relationship between genetic relatedness and microbiota dissimilarity. It is worth re-emphasizing that food and environment was very standardized in this experiment when compared to typical human studies. Genetic relatedness between individuals was measured using genome-wide SNP information while microbiota dissimilarity was measured using Bray-Curtis distance (f.i. Rothschild et al., 2017). We relied on two approaches to mitigate confounding of genetic and environmental effects. In the first we restricted the analyses to full-sibs raised in the same environment, i.e. we confronted genetic similarity and microbiota dissimilarity of litter-mates. In the second we confronted genetic similarity and microbiota dissimilarity across generations (F6 and F7), yet avoiding parent–offspring pairs. Both approaches supported an effect of genetics on microbiota composition manifested by significant negative correlations between genetic similarity and microbiota dissimilarity for (some) individual traits as well as when combining information across traits (Fig. 3A&B). Regressing squared phenotypic difference on genetic distance is an established way to estimate local and global heritability (Haseman & Elston, 1972; Visscher et al., 2006). Yet, Bray-Curtis distance is peculiar in that the phenotypes between which a “difference” is measured are not defined *per se* (Bray and Curtis, 1957). To nevertheless evaluate to what degree of heritability the observed negative correlations might correspond, we simulated quantitative traits with various degrees of heritability in the actual pedigrees and examined the distribution of ensuing correlations between phenotypic distance (absolute value) and genetic distance. These analyses indicated that (in the studied, genetically highly divergent, population) the heritability of microbiota composition may be of the order of ~0.80 within litter, and ~0.20 in the overall population (Suppl. Fig. 7). That the heritability is higher within litter than in the overall population is expected as the environment is obviously more homogeneous within than across litters and generations. Strikingly the impact of genetics was strongest for fecal samples at day 240 in all analyses. This may be in agreement with the observation that, in human, microbiota composition stabilizes with age (Aleman & Valenzano, 2019). Yet, why heritability should be higher in feces than for ileal and cecal content and mucosa remains unclear. Sample types with higher alpha-diversity may be more resilient and hence more heritable.

We also measured the heritability of the abundance of individual taxa. As before, we only extracted within-litter information to mitigate confounding between environment and genetics. Convincing evidence for a genuine influence of host genetics on taxa abundance was the observation of a significant correlation between heritability estimates in the F6 and F7 generation for fecal samples at day 240 and – to a lesser extend – cecum content (Suppl. Fig. 2A). Thus, as for overall microbiota composition, the impact of genetics on abundance of individual taxa appeared highest for feces of mature animals. It is noteworthy that the family with highest heritability in humans (*Christensenellaceae*) also ranked amongst the top raking taxa in the pig data.

Heritability does not accurately foretell the genetic architecture of traits. Phenotypes with low heritability may be affected by variants with major effects (f.i. Kadri et al., 2014), while highly heritable traits may have “omnigenic” architecture (Boyle et al., 2017; Yengo et al., 2018). To gain insight in the genetic architecture of gut microbiota composition in this population we performed GWAS. We identified more than 1,500 signals (corresponding each to a lead SNP x taxon combination) exceeding the genome-wide 5×10^−8^ significance threshold in at least one of the 12 data series. That these include true positive signals was most convincingly demonstrated by the marked shifts towards low p-values when examining the associations between the corresponding lead SNP and taxon in the other data series (Fig. 4A and Suppl. Fig. 3C).

One signal on the telomeric end of chromosome 1 clearly stood out above background noise (experiment-wide significant) in both F6 and F7 cohort, affecting multiple taxa assigned to *Erysipelotrichaceae* (Fig. 4). We showed that this mQTL is caused by a null allele of the ABO gene that results from a 2.3 Kb deletion eliminating 63% of the acetyl-galactosaminyl transferase protein sequence. The corresponding *O* allele was shown to segregate at moderate to high frequency in the eight founder breeds of the mosaic population, in Chinese, Russian and West-European wild boar populations, and in *Sus cebifrons*, a suidae that diverged from the ancestor of the pig ~3.5 million years ago. To gain additional insights in the age of the porcine *O* allele, we generated phylogenetic trees of the A and O alleles of 14 AA and 34 OO animals including domestic pigs, wild boars, Visayan and Javanese warty pigs, and common African warthog. Examination of their local SNP genotypes (50K window encompassing the ABO gene) reveals traces of ancestral recombinations between O and A haplotypes as close as 300 and 800 base pairs from the proximal and distal deletion breakpoints, respectively, as well as multiple instances of homoplasy that may either be due to recombination, gene conversion or recurrent de novo mutations. On their own, these signatures support the old age of the O allele. We constructed UPGMA trees based on nucleotide diversity for windows ranging from 500-bp to 40-Kb centered on the 2.3-Kb deletion. Smaller windows have a higher likelihood to compare the genuine ancestral O versus A states, yet yield less robust trees because they are based on smaller number of variants. Larger windows will increasingly be contaminated with recombinant A-O haplotypes blurring the sought signal. Indeed, for windows ≥ 20-Kb or more, the gene tree corresponds to the species tree, while for windows ≤ 15-Kb the tree sorts animals by AA vs OO genotype (Suppl. Fig. 8). For all windows ≤ 15-Kb the *Sus cebifrons* O allele maps outside of the *Sus scrofa* O allele supporting a deep divergence (rather than hybridization) and hence the old age of the O allele. Of note, for windows ≤1.2-Kb, the warthog A allele is more closely related to the *Sus* A alleles than to the *Sus* O alleles (Suppl. Fig. 8). This suggests that the O allele may be older than the divergence of the *Phacochoerus* and *Sus* A alleles, i.e. > 10 MYA. It will be interesting to study larger numbers of warthog to see whether the same 2.3-Kb deletion exists in this and other related species as well.

This situation in suidae is reminiscent of the trans-species polymorphism of the ABO gene in primates attributed to balancing selection (Ségurel et al., 2012). The phenotype driving balancing selection remain largely unknown yet a tug of war with pathogens is usually invoked: synthesized glycans may affect pathogen adhesion, toxin binding or act as soluble decoys, while naturally occurring antibodies may be protective (Blancher, 2013; Cooling et al., 2015). In humans, the O allele may protect against malaria (Rowe et al., 2007), *E. Coli* and *Salmonella* enteric infection (Robinson et al., 1971), SARS-CoV-1 (Chen et al., 2005), SARS-CoV-2 (Ellinghaus et al., 2020) and schistosomiasis (Camus et al., 1977; Pereira et al., 1979; Ndamba et al., 1997), while being a possible risk factor for cholera (Chaudhuri and De, 1977), *H. pylori* (Boren et al., 1993) and norovirus infection (Lindesmith et al., 2003). Whatever the underlying selective force, it appears to have operated independently in at least two mammalian branches (primates and suidae), over exceedingly long periods of time, and over broad geographic ranges, hence pointing towards its pervasive nature. To gain insights in what selective forces might underpin the observed balanced polymorphism, we tested the effect of ABO genotype on >150 traits measured in the F6 and F7 generations pertaining to carcass composition, growth, meat quality, hematological parameters, disease resistance and behavior. No significant effects were observed when accounting for multiple testing (Suppl. Fig. 9), including those pertaining to immunity and disease resistance.

It is noteworthy that the old age of the “O” allele must have contributed to the remarkable mapping resolution (≤ 3 Kb) that was achieved in this study. In total, 42 variants were in near perfect LD (*r*^2^ ≥ 0.9) with the 2.3 Kb deletion in the F0 generation, spanning 2,298 bp (1,522 on the proximal side, and 762 on the distal side of the 2.3 Kb deletion). This 2.3 Kb span is lower than genome-wide expectations (17th percentile), presumably due to the numerous cross-overs that have accrued since the birth of the 2.3 Kb deletion that occurred in the distant past (Fig. 1E). Yet the number of informative variants within this small segment is higher than genome-wide average of (57% percentile) also probably due at least in part to the accumulation of numerous mutations since the remote time of coalescence of the A and O alleles (Fig. 1D).

The chromosome 1 QTL was the only signal that exceeded experiment-wide discovery and conformation thresholds. QQ-plots obtained after removing chromosome 1 variants (272.8-273.1Mb interval) did not show convincing evidence for residual inflation of log(1/p) values (Suppl. Fig. 3D). This suggests that the residual heritability most likely has a highly polygenic architecture, as becoming increasingly apparent for most complex traits.

The chromosome 1 mQTL was shown to affect the abundance of bacterial species belonging to the family *Erysipelotrichaceae*. The effect was particularly significant for two OTUs (476 and 327) and genus p-75-a5, but affected at least some other *Erysipelotrichaceae* as well. As mentioned above, effects of ABO genotype on host-pathogen interactions are usually interpreted in the context of adhesion or immune response. Yet an alternative mechanism is by altering the source of carbon upon which intestinal bacteria feed. Small and large intestine are amongst the tissues in which the ABO gene is the most strongly expressed (Suppl. Fig. 5A&B). One of its substrates is the heavily glycosylated mucins constituting the intestinal mucus. Mucosal glycans can be used as carbon source by intestinal microorganisms (Mahowald et al., 2009). Glycans first need to be degraded, and the released monosaccharides then imported and catabolized. We reasoned that the mucus of AA/AO pigs would be enriched in GalNAc when compared to OO animals, and that this might favor the growth of bacterial species able to use GalNAc as carbon source. This model makes at least two predictions. The first is that the intestinal GalNAc content should be higher in AA than in OO pigs and this was indeed shown to be the case (Fig. 6A). The second is that the bacteria affected by the mQTL should be able to use GalNAc. We isolated two strains with 16S rRNA sequences that were near-identical to those of the OTU strain (OTU476) that was most affected by the mQTL, and sequenced their complete genome. We showed that it contained the orthologues of eight genes known to be essential for GalNAc import (AgaPTS: *agaE, agaF, agaV, agaW*) and catalysis of the first four GalNAc-specific degradation steps (deacetylation: *nagA*; demanination/isomerisation: *agaS*; kination: *fruK*; aldolase: *gatZ*) hence the five key steps in GalNAc utilization. Importantly, the eight genes clustered in a 50 Kb chromosome segment (Fig. 6C). We generated 3,111 porcine intestinal MAGs from metagenomic shotgun data for comparison. None of these would harbor a GalNAc gene cluster encoding more than four of the five key steps. One catalytic function was always provided in trans, whether GalNAc-6-P deacetylase, tagatose-6-P kinase, or tagatose-1,6-PP aldolase. This finding clearly revealed the unique status of OTU476 with regards to GalNAc utilization. Also consistent with the QTL findings, *Erysipelotrichaceae* MAGs were strongly enriched in clustered GalNAc TR/CP orthologues when compared to MAGs assigned to other bacterial species. Finally, the growth of the two isolated OTU476-like strains was shown to increase when fed with increasing concentrations of GalNAc (Fig. 6D).

Amongst the 3,111 studied MAGs, 15 harbored a gene cluster able to sustain four of the five steps, while the fifth enzyme was encoded somewhere else in the genome. These included one unidentified member of the *Erysipelotrichaceae* family, five strains of *E. Coli*, two *Collinsella* strains (family *Coriobacteriaceae*), two unidentified *Fusobacteriaceae*, and one unidentified *Firmicutes* (Fig. 6C). Gene order within the corresponding GalNAc clusters was supported by observation in two or more independent MAGs and/or by the fact that all concerned genes resided on the same contig. For all 15, the orthologue needed to fulfill the fifth enzymatic reaction (2x tagatose-1-P kinase, 2x GalNAc deacetylase, 1x tagatose-1,6-PP aldolase) was found somewhere else in the genome, allowing us to assume that all these species are able to utilize GalNAc. Fifty additional MAGs contained the orthologues needed to accomplish the five key steps in GalNAc utilization albeit without evidence for a similar degree of clustering (either because the genes are indeed not clustered in the corresponding genomes or because they were segregated across distinct sequence contigs). It is reasonable to assume that several of those bacteria are also able to utilize GalNAc as carbon source. Why then would the chromosome 1 mQTL only affect a small subset of *Erysipelotrichaceae* species? We first reasoned that OTU476-like strains might be more dependent on GalNAc availability than other species, for instance because they can’t utilize alternative, common monosaccharides as carbon source. To test this hypothesis, we searched for KEGG numbers that would commonly occur in other genomes, yet were absent in the OTU476-like strains. We performed this analysis for all MAGs, as well as separately for the MAGs that were predicted to be able to use GalNAc (cfr. above). There was no convincing evidence that OTU476 might be missing a common and important monosaccharide-utilizing pathway in their genome (data not shown). Closer examination of the structure of the most complete GalNAc gene clusters in the studied MAGs (Fig. 6C) revealed an alternative, possible clue. The GalNAc gene clusters of the *non-Erysipelotrichaceae* species all have the features expected from genuine regulons. The relevant ORFs tend to be adjacent to each other (spanning ~ 10Kb) and on the same strand, hence compatible with poly-cistronic messenger RNAs enabling coregulated expression. In striking contrast, the ORFs of the GalNAc clusters of the OTU476-like strains and at least one studied *Erysipelotrichaceae* are spanning respectively ~50 and ~30Kb, and appear to be distributed randomly on both strands. Most importantly, neither genome contained orthologues of *agaR* (K02081) or *gntR* (K03710), which are encoding negative regulators of GalNAc regulons and were observed in all other GalNAc-rich MAGs. It is noteworthy that out of the 77 *Erysipelotrichaceae* MAGs encompassing an orthologue of *agaS*, only two (=2.6%) had an orthologue of *agaR* or *gntR* in its vicinity. This number has to be compared with the fact that out of the 85 “Other” (i.e. non-*Erysipelotrichaceae*) MAGs encompassing an orthologue of *agaS*, 39 (=46.4%) had such *agaR* or *gntR* orthologue in *agaS’s* vicinity. The *fruR* repressor that is observed in the vicinity of the GalNAc genes in the OTU476-like strains was found in the vicinity of *agaS* in only 1/77 instances in *Erysipelotrichaceae* and 3/85 instances in other MAGs, indicating that the *FruR-AgaS* colocalization in OTU476-like strains is likely coincidental. Taken together, this suggests that, contrary to *E. Coli* and other bacterial species, the OTU476-like strains and some *Erysipelotrichaceae* are not endowed with the capacity to sense GalNAc concentrations in the medium and only induce expression of the genes and proteins necessary for GalNAc utilization when needed (Leyn et al., 2012; Biddart et al., 2014; Zhang et al., 2015), but may rather express their GalNAc-related genes constitutively. The GalNAc gene cluster as seen in the OTU476-like strains is a possible evolutionary intermediate towards the formation of a genuine regulon as seen in *E.Coli*, already facilitating horizontal transmission of a “selfish” functional gene ensemble even if not yet adaptively coregulated (Lawrence, 1999). This testable hypothesis (constitutive versus inducible expression) suggests an alternative modus operandi of the chromosome 1 QTL. Against intuition, bacteria affected by the mQTL (i.e. OTU-476, OTU-327, p-75-a5 and some other *Erysipelotrichaceae*) may very well not be at an advantage when GalNAc is present at high concentration in the intestinal content (as in AA and AO animals), but rather at a disadvantage when GalNAc is present at low concentrations (as in OO animals) because then they waste energy transcribing and translating useless genes. By regulating expression of their GalNAc operon in response to ambient GalNAc availability, species like *E. Coli* may fair equally well in the gut of AA/AO as in that of OO pigs, hence not be affected by the mQTL. It is worth noting that the A allele is dominant with regards to OTU476, OTU327 and p-75-a5 abundance (Fig. 5B), suggesting that the additional increase in GalNAc concentrations in AA (vs AO) animals does not further benefit these taxa.

We examined the effect of ABO blood group on the abundance of ~75 OTUs assigned to *Erysipelotrichaceae* in human gut samples. Although we could not rigorously test this for all OTUs (as some human V1-V2 and V5-V6 data could not directly be compared with porcine V3-V4 data) none of the OTUs detected in human samples were as closely related to the pig OTU-476, OTU-327 or p75.a5 as these were to each other. We found no evidence for an effect of ABO blood group on the abundance of any of these OTUs. What underlies the difference between pigs and humans is unclear. Either strains susceptible to ABO genotype are not present at sufficient frequency in human feces, or the carbohydrate composition of human intestinal content makes these strains less sensitive to variations in GalNAc concentrations. It is worth noting that the studied human samples were intestinal biopsies collected after a standard gut cleansing procedure. The abundance of the genus p-75-a5 was recently found to differ significantly between African subsistence categories and to be highest in pastoralists (as compared to hunter-gatherers and agro-pastoralists) possibly as a result of interaction with livestock (Malmuthuge et al., 2014; Hansen et al., 2019). Repeating the experiments in pastoralist populations may reveal the same mQTL effect detected in this study.

**Supplemental Figure 7:**
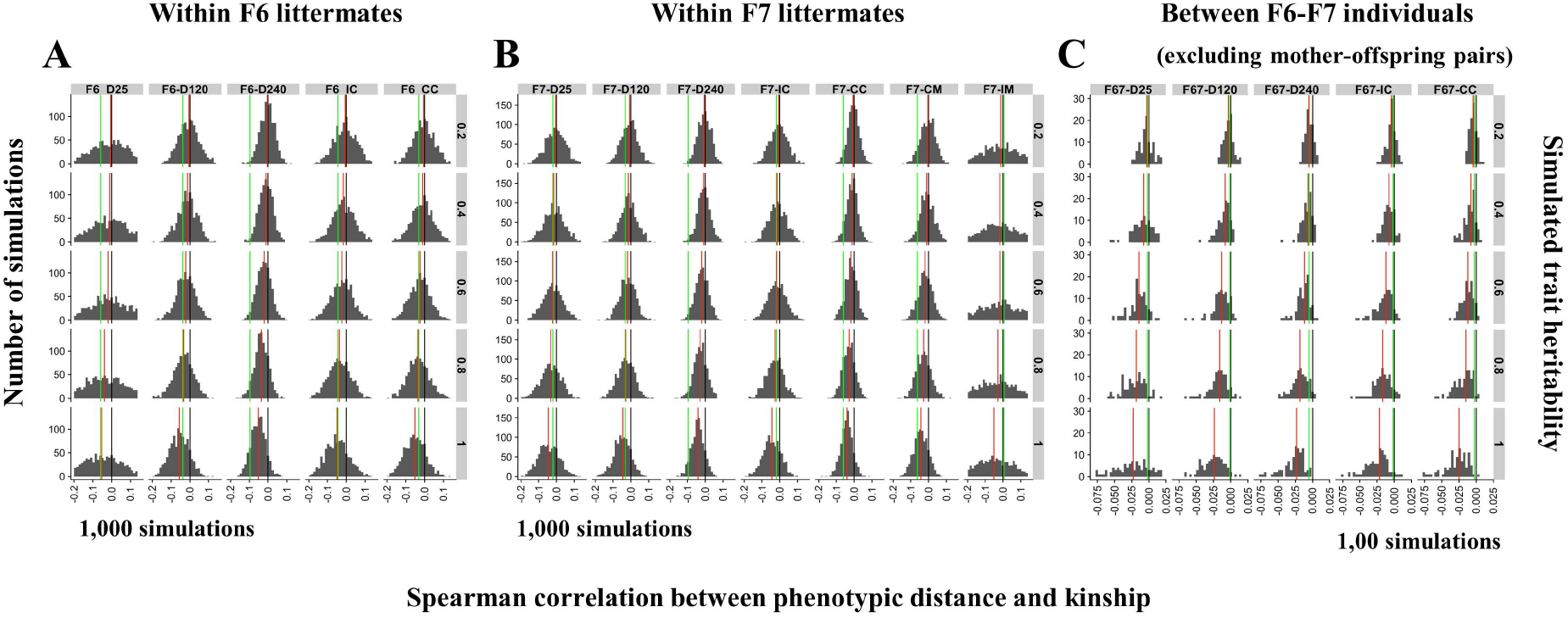
We observed an excess of negative correlations between genetic similarity (from SNP genotype data) and microbiota dissimilarity (Bray Curtis distance computed from 16S rRNA data) both within litter as well as between generations, supporting an effect of host genetics and intestinal microbiota composition (Fig. 3A&B). We took care in these analyses to mitigate effects of litter on both genetic and microbiota distance metrics, (as these may inflate statistical significance) by applying permutations tests. Regressing squared phenotypic difference on genetic distance is a standard way to estimate local or global heritability (Haseman & Elston, 1972; Visscher et al., 2006). It can be shown that 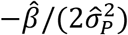 estimates the narrow sense heritability 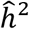. In these, 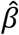 is the least square regression coefficient and 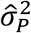 an estimate of the phenotypic variance. In our analyses, and following standard procedures (f.i. Rothschild et al., 2017), we used Bray-Curtis as distance measure for microbiota composition. For this metric, there is no corresponding individual phenotype *p_i_ per se*. We therefore used simulations to translate the observed negative correlations in measures of heritability. For the within-generation/within-litter analyses, we first used the actual measures of kinship for all litter-mates computed with GEMMA (Zhou & Stephens, 2012) and corresponding to the x-axis in Fig. 3A. We standardized them (mean 0 and SD 1), scaled them (mean of 0.5 and SD 0.04, following Visscher et al., 2006) and multiplied them by *h*^2^ (0.2, 0.4, 0.6, 0.8 or 1.0). We sampled “breeding values” from a multivariate normal distribution with means 0 and corresponding variance-covariance matrix using the *mvrnorm* R function. For each individual, we sampled an environmental effect from a normal distribution with mean 0 and variance (1 – *h*^2^) using the *rnorm* R function. Breeding values and environmental effects were added to yield a phenotypic value *p_i_* for each individual. We then computed Spearman’s rank correlation between *abs*(*p_i_* – *p_j_*) and Θ_*ij*_ for all pairs of litter mates *i* and *j* using the *cor.test(method=”spearman”)* R function. In this Θ_*ij*_ is the kinship metric computed by GEMMA. We repeated the simulations 1,000 times. Suppl. Fig. 7A&B show the distribution of the corresponding correlations (*r*) for the 12 data series and 5 values of *h*^2^. The black vertical line corresponds to zero. The red vertical line to the median of the simulations. It can be seen that as the heritability increases the value of the median decreases as expected. The green lines correspond to the corrected correlation (*r_c_*) obtained with the real data. A rough estimate of the heritability of the real trait (microbiota composition) was deduced from the coincidence between the red and green lines. As an example, the heritability of microbiome composition for data series F7-D120 was assumed to be close to 0.8. We proceeded in the same way for the across generation analysis (Suppl. Fig. 7C). We used the actual measures of kinship across the F6 and F7 generations computed with GEMMA. We standardized them (mean 0 and SD 1), and then scaled them such that the values for an individual with itself would center on 1, and for full-sibs on 0.5. Breeding values and environmental effects were sampled using *mvrnorm* and *rnorm* as above. As the number of individuals is much higher in these analyses we only performed 100 simulations.

**Supplemental Figure 8:**
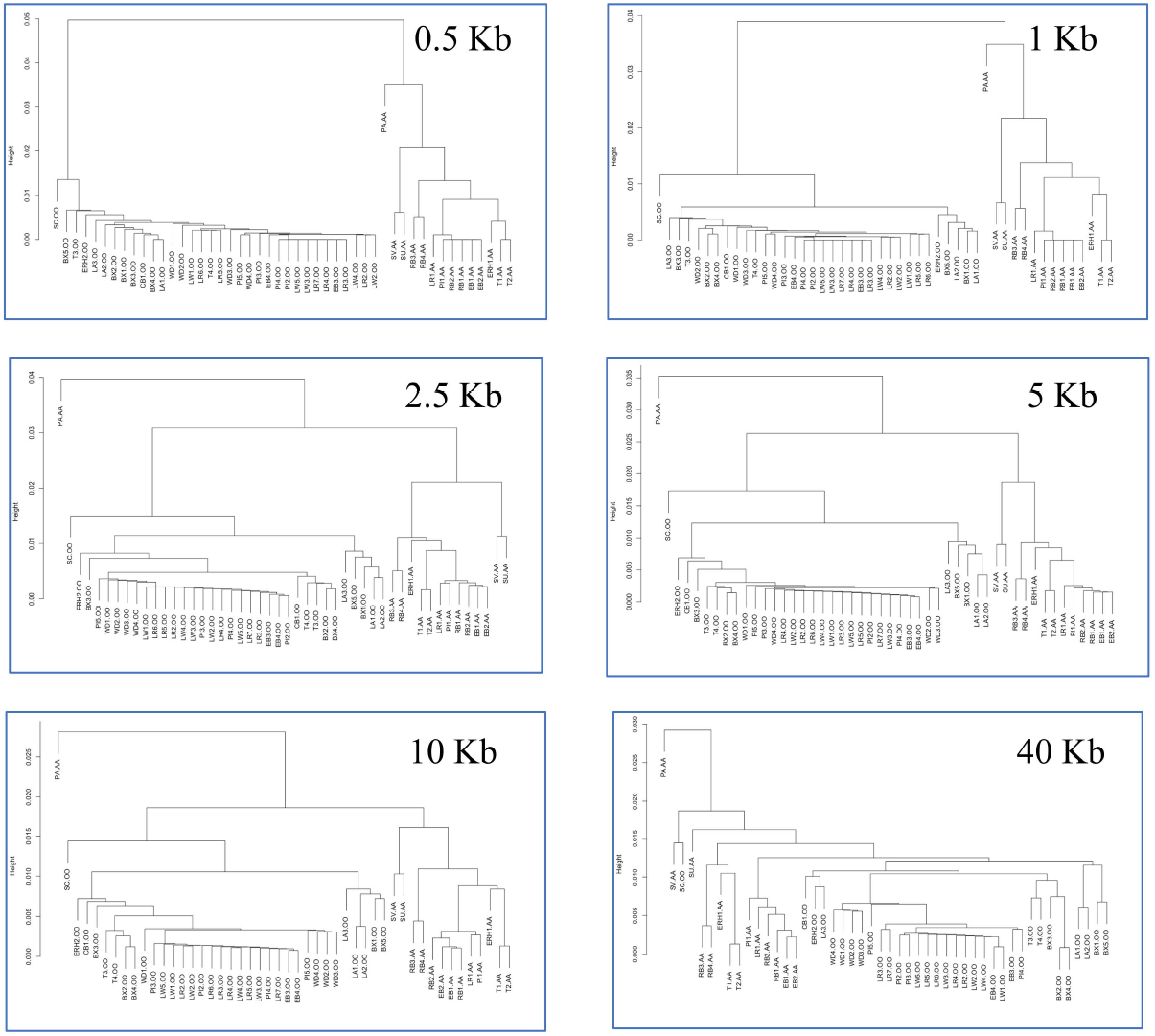
UPGMA tree based on nucleotide diversities between 14 AA and 34 OO animals in windows of increasing size (0.5 to 40-Kb) centered on the 2.3 Kb deletion in the ABO gene (porcine O allele). PA: *Phacochaerus Africanus*, SC: *Sus cebifrons*, SV: *Sus verrucosus*, SU: *Sus scrofa vittatus*, CB: Chinese wild boar, RB: Russian wild boar, EB: European wild boar, ERH: Erhualian, BX: Bamaxiang, T: Tibetan, LA: Laiwu, LR: Landrace, LW: Large White, PI: Piétrain, WD: White Duroc.

**Supplemental Figure 9:**
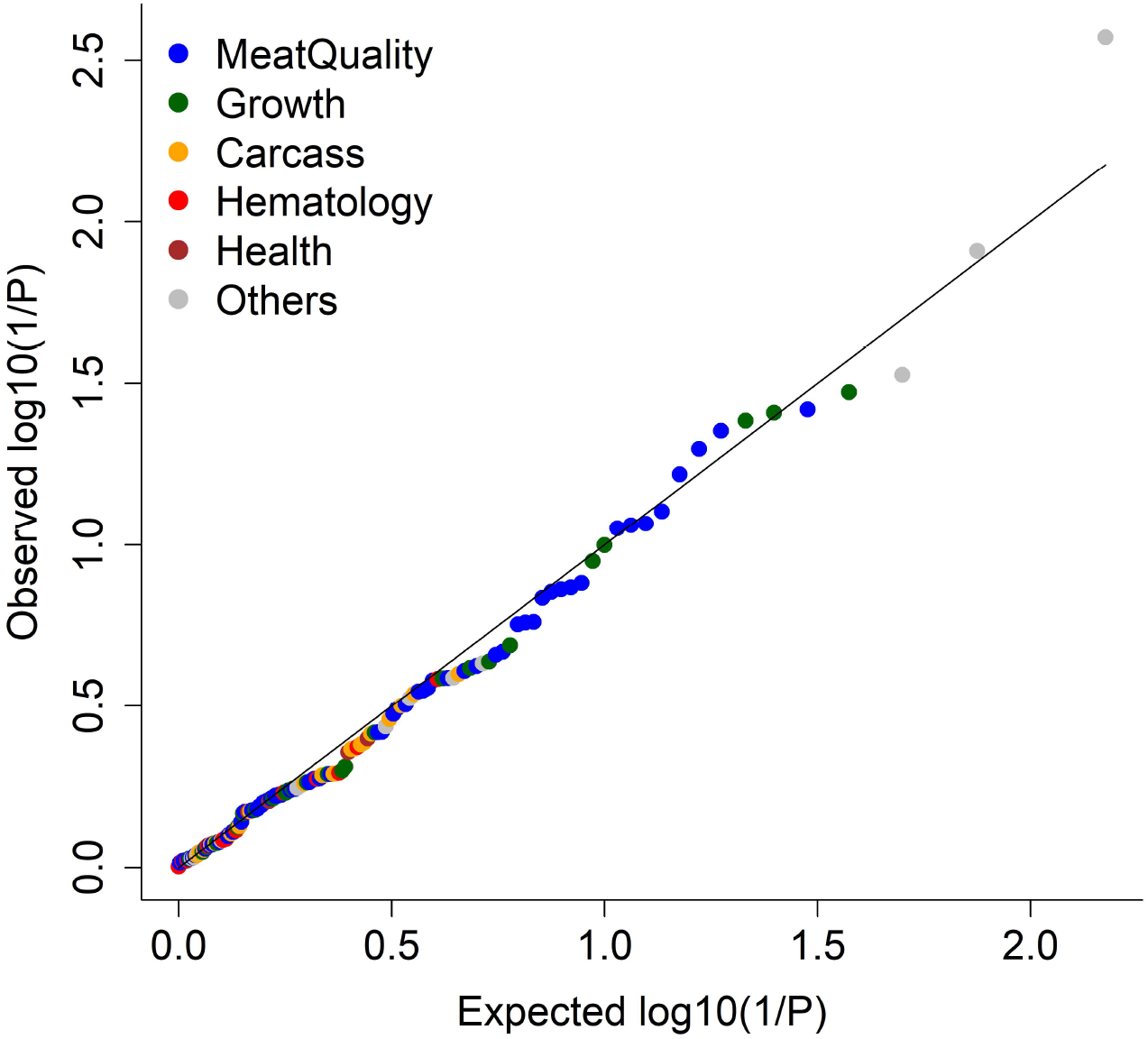
QQ plots for the effect of AO genotype on 150 phenotypes pertaining to meat quality, growth, carcass composition, hematology, health, and other phenotypes in the F6 and F7 generation. The p-values were obtained by meta-analysis (weighted Z score) across the F6 and F7 generations.

## STAR METHODS

### Animal rearing and sample collection

This study focused on the sixth (F6) and seventh (F7) generation of a mosaic population generated as follows. An average of 3.6 boars (range: 3 – 4) and 4 sows (range: 2 – 5) from four indigenous Chinese pig breeds (Erhualian (EH), Bamaxiang (BX), Tibetan (TB), Laiwu (LA)) and four commercial European/American pig breeds (Landrace (LD), Large White (LW), Duroc (WD) and Piétrain (PT)) were successfully applied in the mating design, thus, constituted the F0 generation. For each Chinese breed, the boars were mated with the ewes of one European breed, and the sows with the boars of another European breed to produce the F1 generation. Thus, every Chinese and every European breed is parent breed of two distinct F1 hybrid combinations each, for a total of eight F1 combinations (BX-LW, BX-PT, LA-PT, LA-LD, TB-LD, TB-WD, EH-WD, EH-LW). The F2 generation was obtained by mating each F1 hybrid combination with two others that did not share parental breeds for a total of eight F2 combinations (BX-LW x LA-PT, BX-PT x LA-LD, LA-PT x TB-LD, LA-LD x TB-WD, TB-LD x EH-WD, TB-WD x EH-LW, EH-WD x BX-LW, EH-LW x BX-PT). Every F2 combination was obtained by reciprocally crossing an average of 4 boars from one F1 combination with an average of 7.25 sows from the other. The F3 generation was obtained by mating each of the eight F2 hybrid combinations with the only complementary F2 combination that did not share any parental breeds for a total of four F3 combinations (BX-LW-LA-PT x TB-LD-EH-WD, BX-PT-LA-LD x TB-WD-EH-LW, LA-PT-TB-LD x EH-WD-BX-LW, LA-LD-TW-WD x EH-LW-BX-PT) expected to each have ~12.5% of their genome from each of the founder breeds. Every F3 combination was obtained by reciprocally crossing an average of 7 boars from one F2 combination with an average of 10.8 sows from the complementary one. The F4, F5, F6 and F7 generations were obtained by intercrossing 57 boars x 75 sows (F3->F4), 62 boars x 97 sows (F4->F5), 85 boars x 170 sows (F5->F6), and 82 boars x 111 sows (F6->F7)(Suppl. Table 1).

All F6 and F7 animals were born and reared at the experimental farm of the National Key Laboratory for swine Genetic Improvement and Production Technology, Jiangxi Agricultural University (Nanchang, Jiangxi) under standard and uniform housing and feeding conditions. Piglets remained with their mother during the suckling period and were weaned at ~46 days of age. Litters were transferred to 12-pig fattening pens with automatic feeders (Osborne Industries, US), minimizing splitting and merging of litters. All pigs were fed twice per day with formula diets containing 16% crude protein, 3,100 kJ digestible energy, 0.78% lysine, 0.6% calcium and 0.5% phosphorus. Water was available ad libitum from nipple drinkers. Males were castrated at 80 days. Fecal samples were manually collected from the rectum of experimental pigs at the ages of 25, 120 and 240 days, dispensed in 2ml tubes, flash frozen in liquid nitrogen, and stored at −80°C. Animals were slaughtered at day 240. Ileum and cecum were sealed at both ends with a sterile rope and extracted from the carcass. Within 30 min after slaughter, ileal and cecal luminal content were collected (F6 and F7 animals), ileum and cecum rinsed with sterile saline solution, and samples of ileal and cecal mucosa scraped with a sterile microscopic slide (F7 animals only). Approximately one gram of content or scrapings was packed in 2-ml sterile freezer tubes, flash frozen in liquid nitrogen, and stored at −80°C. The number of samples of the different types available for further analysis are provided in Suppl. Table 2. All the animals included in the analyses were healthy and did not receive any antibiotic treatment within one month of sample collection. All procedures involving animals were carried out according to the guidelines for the care and use of experimental animals established by the Ministry of Agricultural and Rural Affairs and Jiangxi Agricultural University.

### Genotyping by sequencing of the F0, F6 and F7 generations

Genomic DNA was extracted from ear punches using a standard phenol-chloroform-based DNA extraction protocol. DNA concentrations were measured using a Nanodrop-1000 instrument (Thermo Scientific, USA), and DNA quality of all samples assessed by agarose (0.8%) gel electrophoresis. Genomic DNA was sheared to 300-400 bp fragment size. 3’-ends were adenylated and indexed primers ligated. Libraries were amplified by PCR using Phusion High-Fidelity DNA polymerase (NEB, USA) following the recommendations of the manufacturer (Illumina, US). The libraries were loaded on Illumina X-10 instruments (Illumina Inc., San Diego, CA) for 2 × 150 bp paired-end sequencing by Novogene (Beijing, China). We removed reads with quality score ≤ 20 for ≥50% of bases or ≥ 10% missing (=”N”) bases. Read quality was checked using Fastqc (http://www.bioinformatics.babraham.ac.uk/projects/fastqc/). Clean reads were aligned to the Sus scrofa reference genome assembly 11.1 (Warr et al., 2019) using BWA (Li & Durbin, 2010). Bam files of mapped reads were sorted by chromosome position using SAMTools (Li et al., 2009). Indel realignment and marking of duplicates were done with Picard (http://broadinstitute.github.io/picard). Individual genotypes were called from BAM files using Platypus (v0.8.1) (Rimmer et al., 2014). Individual genotypes were merged into a single VCF file using PLINK (v1.9) (Chang et al., 2015) encompassing a total of 39.3 million variants including 31,094,663 SNPs and 8,266,390 INDELs. Missing genotypes were imputed with Beagle (v.40) (Browning & Browning, 2007). Genomic variants with minor allele frequencies (MAF) < 0.03 were removed.

### Computing nucleotide diversities

Nucleotide diversities between pairs of breeds were computed from variant frequencies as follows:

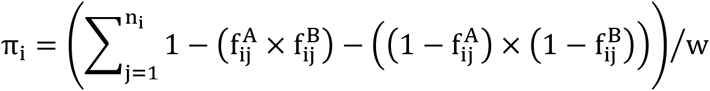

where π_i_ is the nucleotide diversity in window i, n_i_ is the number of variants in window i, 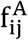 is the frequency of variant j of window i in breed A, 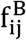 is the frequency of variant j of window i in breed B, and w is the size of the windows in base pairs. The overall nucleotide diversity for a pair of breeds A and B was computed as the average of π_i_ across all windows. The numbers reported are averages of overall nucleotide diversities for multiple pairs of breeds (within European, within Chinese, between European, between Chinese, between European and Chinese), computed for a window size of 1 million base pairs.

### Estimating the contribution of the eight founder breeds in the F6 and F7 generation at genome and chromosome level

We estimated the proportion of the genome of the eight founder breeds in the F6 and F7 generation following Coppieters et al. (2020). Assume that the total number of variants segregating in the mosaic population is n_T_. Each of these variants has a frequency in each one of the founder breeds which we denote 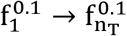 for breed 1, 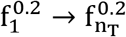 for breed 2, etc … as well as a frequency in the F6 (or F7) generation which we refer to as 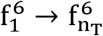. We assume that there is a total of B breeds. We denote the proportion of the genome of breed 1 in generation F6 (or F7) as P_1_, of breed 2 in generation F6 (or F7) as P_2_, etc … We estimated the values of P_1_, P_2_, etc. … using a set of linear equations:

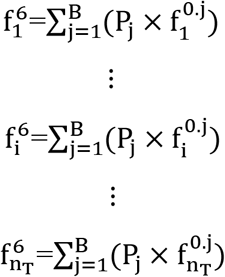

We used standard least square methods (lm function in R) to find the solutions of P_j_ that minimize the residual sum of squares. This was done for the entire genome, as well as by autosome.

### 16S rRNA data collection and processing

Microbial DNA was extracted from feces, luminal content and mucosal scrapings using the QIAamp Fast DNA stool Mini Kit following the manufacturer’s recommendations (Qiagen, Germany). DNA concentrations were measured using a Nanodrop-1000 instrument (Thermo Scientific, USA), and DNA quality assessed by agarose (0.8%) gel electrophoresis. The V3 – V4 hypervariable region of the 16S rRNA gene was amplified with the barcode fusion primers (338F: 5-ACTCCTACGGGAGGCAGCAG-3, 806R: 5-GGACTACHVGGGTWTCTAAT-3) with 56 °C annealing temperature. After purification, PCR products were used for constructing libraries and sequenced on an Illumina MiSeq platform (Illumina, USA) at Major bio (Shanghai, China). The 16S rRNA sequencing data were submitted to the CNGB database and have accession number CNP0001069. The raw 16S rRNA gene sequencing reads were demultiplexed and primer and barcode sequences trimmed using Trimmomatic (V.0.39) (Bolger et al., 2014). Reads with ≥10 consecutive same or ambiguous bases were eliminated. Clean paired-end reads were merged (minimum 10 bp overlap) into tags using FLASH (v.1.2.11) (Magoc & Salzberg, 2011). The average number of tags per sample was ~ 40,888 (Suppl. Table 2). Chimeric reads were removed using USEARCH (v.7.0.1090) (Edgar, 2010). Sequence data were rarefied to 19,631 tags, i.e. the lowest number of tags per sample. Tags were clustered in operational taxonomic units (OTUs) with VSEARCH (v.2.8.1) (Rognes et al., 2016) using 97% similarity threshold. OTUs that would not have ≥ 3 reads in at least two samples or were detected in ≤ 0.2% of the samples were ignored. In the end, 12,054 OTUs accounting for an average of 98.7% of total reads per sample were used for further analysis. OTUs were matched to taxa using the Greengenes (v13.5) database and the RDP classifier (v2.2) (Wang et al., 2007). Principal coordinate analysis (PCoA) was performed with the “ape” and “vegan” R packages using Bray-Curtis dissimilarities. Shannon’s index was used as α-diversity metric and computed using mothur (v 1.43.0) (Schloss et al., 2009). Bray-Curtis dissimilarity was used as β-diversity metric and computed using vegdist of the vegan package in R. The mouse fecal microbiome data were from Cheema et al. (2019). The human fecal microbiome data were 16S rRNA data from 106 healthy individuals (Shagam et al., in preparation).

### Measuring the heritability of microbiome composition

We first estimated the impact of host genetics on the composition of the intestinal microbiome by measuring the correlation between genome-wide kinship and microbiome dissimilarity. We computed genome-wide kinship (Θ) for all pairs of relevant individuals (see hereafter) using the SNP genotypes at the above-mentioned 30.2 million DNA variants using either GEMMA (Zhou & Stephens, 2012) or GCTA (Yang et al., 2011). Both programs yielded estimates of Θ with same distribution after standardization, albeit different raw values. We herein report results obtained with GEMMA. Microbiome dissimilarity was measured using the Bray-Curtis dissimilarity computed using the “vegan” R function (Dixon, 2003) and abundances of all OTUs. We computed Spearman’s (rank-based) correlations using the “corrtest” function in R (v3.5.3). We first performed this analysis for each trait and generation separately within litter, i.e. only considering pairs of full-sibs born within the same litter, hence in essence following Visscher et al. (2006). We then performed the analysis across the F6 and F7 generations. The pairs of individuals considered were all F6-F7 animal pairs except sow-offspring. To account for dependencies characterizing the data the statistical significance of the obtained correlations was determined empirically by permutation testing (1,000 permutations): vectors of OTU abundances were permuted within litters, Bray-Curtis distances recomputed, and correlated with the unpermuted kinships. The empirical p-value was determined as the proportion of permutations that yielded a Spearman’s correlation that was as low or lower than that obtained with the real data. Spearman’s correlation coefficients were then adjusted to match the empirical p-values as follows. We generated “breeding values” for all animals used in Fig. 3 by sampling from a multivariate normal distribution which variance corresponding to the simulated heritability (h^2^) and covariance matrix constrained by the actual pairwise kinship coefficients using the mvrnorm R function. We added environmental effects to the breeding values (yielding phenotypic values) sampled from a normal distribution with mean 0 and variance (1 – h^2^) using the rnorm R function. We then computed the corresponding Spearman’s correlation between the pairwise genetic distances and the absolute value of the pairwise phenotypic differences using the cor.test(method=”spearman”) R function. The corrected Spearman’s correlation was then chosen as the one obtained with the simulated data set (out of 5,000) that yielded a one-sided p-values that was the closest one to the p-value obtained by permutation with the corresponding real data set. See also legend to Supplemental Figure 7.

Heritabilities of uncorrected abundances of specific taxa were estimated using a linear mixed model implemented with GEMMA (Zhou & Stephens, 2012). The model included a random polygenic and error effect and no fixed effects. Variance components were estimated with GEMMA. Analyses were conducted separately for the 12 data series. To obtain unbiased estimates of h^2^, we repeated the analysis 1,000 times after permutation of the taxa abundances within litter. The average h^2^ obtained across permuted datasets was subtracted from the h^2^ obtained with the real (i.e. unpermuted) data to yield an unbiased estimate 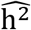. The statistical significance of 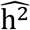 was estimated as the proportion of permutations that would yield a value of h^2^ that would be as high or higher than the value of h^2^ obtained with the real data. To provide further support for the validity of the h^2^ estimates, we measured Spearman’s correlation between F6 and F7 estimates (both h^2^ and corresponding −log(p) values) computed with the corr.test R function. The “total heritability” of the intestinal microbiome was further computed from the heritabilities of individual taxa abundance following Rothschild et al. (2018).

### Mapping microbiota QTL (mQTL)

mQTL were mapped using the GenABEL R package (Aulchenko et al., 2007) applying two models following Turpin et al. (2016). The first fitted a linear regression between allelic dosage and log_10_-transformed taxa abundance. It was applied to all SNPs with MAF ≥ 0.05 (in the corresponding data series) and taxa with non-null abundance in at least 20% of samples (in the corresponding data series), ignoring samples with null abundance if those represented more than 5% of samples. The second fitted a logistic regression model between allelic dosage and taxon presence/absence in the corresponding sample (binary model). It was applied to all SNPs with MAF ≥ 10% (in the corresponding data series) and taxa present in ≥ 20% and ≤ 95% of samples (in the corresponding data series). Both models included sex, slaughter batch (21 for F6, 23 for F7) and the three first genomic principal components as fixed covariates. GWAS were conducted separately for each taxon x data series combination and p-values concomitantly adjusted for residual stratification by genomic control. P-values were combined across traits and/or taxa using a z-score. P-values were converted to signed z-values using the inverse of the standard normal distribution and summed to give a “z-score”. Z-scores were initially calculated using METAL (Willer et al., 2010). To compute the p-value of the corresponding Z-score while accounting for the correlation that exists between the phenotypic values of a given cohort across traits we also computed the genome-wide (i.e. across all tested SNPs) average 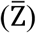 and standard deviation (σ_Z_) of the Z score. The p-value of Z scores was (conservatively) computed by assuming that 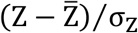 is distributed as N(0,1) under the null hypothesis. Both approaches yielded similar results.

### De novo assembly of the A allele of the porcine ABO acetyl-galactosaminyl transferase gene

We extracted high-quality genomic DNA from longissimus dorsi of a Bamaxiang female using a phenol-chloroform-based extraction method (Novogene Biotech, Beijing, China). A 40 kb SMRTbell DNA library (Pacific Biosciences of California, CA, USA) was prepared using BluePippin for DNA size selection (Sage Science, MA, USA) and then sequenced on a PacBio Sequel platform (Pacific Biosciences of California, CA, USA) with P6/C4 chemistry at Novogene Biotech, Beijing, China. We obtained a total of 18,148,470 subreads with N50 length of 17,273 bp. Additionally, a paired-end library with insert size of 350-bp was constructed and sequenced on an Illumina Novaseq 6000 PE150 platform (2×150bp reads) at Novogene Biotech, Beijing, China. PacBio reads were self-corrected using Canu (v1.7.1) before assembly with Flye (v2.4.2) (Kolmogorov et al., 2019). Errors in the primary assembly were first corrected using PacBio subreads using racon (v1.4.10)(Vaser et al., 2017), and Illumina paired-end reads were then mapped to the contigs using bwa-mem (Li, 2013) to polish the contigs using Pilon (v1.23, Broad Institute, MA, USA) (Walker et al., 2014). Lastz (Harris, 2007) and Minimap2 (v2.17-r941) (Li, 2018) were used to compare the Bamaxiang contig and the 40k sequence spanning the ABO gene of the Sus scrofa Build 11.1 reference genome.

### Developing a PCR assay to distinguish AA, AO and OO pigs

We designed two pairs of primers to genotype the deletion in the F6 and F7 populations. The first pair of primers was located within intron 7 of the ABO gene and downstream of the deletion (FP: 5’-GAGTTCCCCTTGTGGCTCAGT-3’, RP: 5’-TTGCCTAAGTCTACCCCTGTGC-3’). The second pair of primers was located in exon 8 (FP2: 5’-CGCCAGTCCTTCACCTACGAAC-3’, RP2: 5’-CGGTTCCGAATCTCTGCGTG-3’). PCR amplification was performed in a 25-μl reaction containing 50 ng genomic DNA and 1.5 U of LA Taq DNA polymerase (Takara, Japan) under thermocycle conditions of 94°C for 4 minutes, 35 × (94°C for 1 min, 1 min at specific annealing temperature for each set of primers and 72°C for 2 min), and 72°C for 10 minutes on a PE 9700 thermal cycler (Applied Biosystem, USA).

### RNA seq and eQTL analysis

A total of 300 cecum tissue samples from F7 pigs which also had microbiota and genotype data were used to extract total RNA with TRIzol™ (Invitrogen, USA) following the manual. Total RNA was electrophoresed on 1% agarose gel. RNA purity and integrity were assessed using an eNanoPhotometer^®^ spectrophotometer (IMPLEN, USA) and a Bioanalyzer 2100 system (Agilent Technologies, USA). Qubit3.0 Fluorometer was used to measure RNA concentration. 2-μg total RNA of each sample were used to constructed RNA sequencing libraries, using the NEBNext^®^ UltraTMR NA Library Prep Kit for Illumina (NEB, USA) following the manufacturer’s protocol. Briefly, Oligo (dT) magnetic beads (Invitrogen, USA) were used to enrich mRNA, which was then fragmented using a fragmentation buffer (Ambion, USA). cDNA was synthesized by using 6-bp random primers and reverse transcriptase (Invitrogen, USA). After purification, cDNA was end-repaired, and index codes and sequencing adaptors ligated. After PCR amplification, purification and quantitation, the libraries were sequenced on a Novaseq-6000 platform using 2×150-bp paired-end sequencing. Clean data were obtained by removing adapter reads, poly-N and low-quality reads from raw data. Cleaned reads from each sample were mapped to the complete ABO sequence from the Bamaxiang reference genome with the A allele at the ABO locus constructed by the authors using STAR (Dobin et al., 2013). Samtools (Li et al., 2009) was used to convert SAM format to BAM format. The read counts mapping to ABO (exon 1 to 7) were quantified for each sample using featureCounts (Liao et al., 2014). To adjust for the effect of sequencing depth, the expression abundance of ABO gene was normalized to fragments per kilobase of exon model per million mapped reads (FPKM). Gender and batch were treated as covariates to correct for gene expression levels, and the corrected residuals used for subsequent analyses. GEMMA (Zhou & Stephens, 2012) was used to analyze the association of ABO expression level with genome-wide variants using a linear mixed model.

### Whole-genome sequencing and bioinformatic analysis for wild boars, Sus verrucosus and Sus cebifrons

The genomes of six Russian wild boars, one Sumatran wild boar, and one African warty hog were sequenced on an Illumina HiSeq X Ten platform at Novogene Biotech, Beijing, China. Additionally, six Chinese wild boars were sequenced in a previous study (Ai et al. 2015), and we downloaded the genome sequence data for eight other pigs from NCBI. Finally, we used a total of 22 genomes to call SNPs in the porcine ABO gene using GATK (Van der Auwera et al., 2013). We replaced the ABO gene of the Sus scrofa build 11.1 genome with the 50 Kb Bamaxiang contig sequence containing the A allele of ABO gene. The cleaned reads of the 22 individuals were aligned to the modified Sus scrofa reference genome (build 11.1) using BWA (Li and Durbin, 2010).

### Phylogenetic analysis of the O alleles in the Sus genus

We applied GATK to perform indel realignment, and proceeded to SNP and INDEL discovery and genotyping with UnifiedGenotyper across all 83 samples simultaneously using standard hard filtering parameters according to GATK Best Practices recommendations (DePristo et al., 2011). We restricted the analysis to the 14 AA and 34 OO animals (Fig. 5C), hence circumventing the need to phase the corresponding genotypes. We defined windows of varying size (0.5 to 50Kb) centered around the 2.3 Kb deletion. For all pairs of individuals, we computed a running sum over all variants in the window adding 0 when both animals had genotype AA_ (alternate) or RR(reference), 1 when one animal was AA and the other RR, and 0.5 in all other cases. The nucleotide diversity for the corresponding animal pair was then computed as the running sum divided by the window size in bp. We ignored the variants located in the 2.3 Kb deletion in this computation. The ensuing matrix of pair-wise nucleotide diversities was then used for hierarchical clustering and dendrogram construction using the hclust(method=”average”) R function corresponding to the unweighted pair group method with arithmetic mean (UPGMA).

### Analysis of population differentiation

We quantified the degree of population differentiation by computing the effect of breed on the variance of allelic dosage using a standard one-way ANOVA fixed effect model and a F-statistic computed as the ratio of the “between breed mean squares” (BMS) and “within breed mean squares” (WMS) (Weir & Cockerham, 1984). BMS and WMS were computed as:

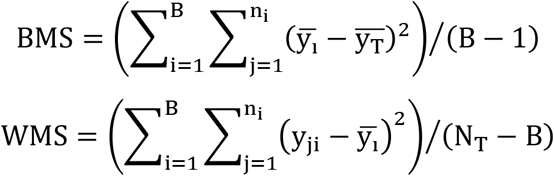

where y_ij_ is the allelic dosage of the alternate allele in individual j (of n_i_) of breed i (of B), 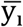 is the average allelic dosage in breed i (of B), and 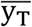 is the average allelic dosage in the entire data set. We computed the average of the corresponding F-statistic for all variants within a sliding window of fixed physical size (f.i. 2-Kb in Fig. 5E), and took the inverse of this mean as measure of population “similarity”. The corresponding profiles were nearly identical to those obtained by computing average F_ST_ values across variants (and taking the inverse) following Nei (1977).

### Determination of the concentration of N-acetyl-galactosamine in cecal lumen

Targeted LC-MS/MS analysis was performed to determine the concentration of N-acetyl-galactosamine in cecal lumen samples using a liquid chromatography mass spectrometry system comprising an ExionLC™ AD System (AB Sciex, USA) coupled to a TripleTOF™ 5600 Mass Spectrometer (AB Sciex, USA). Cecal lumen samples used for measurement of N-acetyl-galactosamine were harvested from F7 pigs which had microbial composition data and were used for GWAS. The samples were thawed from −80°C. Approximately 0.2g of each sample was homogenized in double-distilled water and centrifuged (5000 r.p.m., 3min; 12000 r.p.m., 20min, at 4°C). The supernatants were filtered and submitted to measurement on the LC-MS/MS instrument. Separation was performed in a 2.1 x 100mm, 1.7um ACQUITY UPLC BEH C18 Column (Waters, USA). DuoSpray-MS/MS was performed in positive ion mode with two scan events: MS and MS/MS scan with mass range of m/z 100-1000 and 100-250, respectively. In the product ion, the ionspray voltage was set at 5.5kV, the temperature was maintained at 500°C, and the collision energies were optimized from ramping experiments. In addition, the standard substance of N-acetylgalactosamine (Aladdin, China) solutions were applied to LC-MS/MS to determine the peak elution time and m/z value of precursor ion and product ion as a reference.

### Isolating 4-8-110 and 4-15-1

Fecal samples were collected from the rectum of healthy pigs at about 120 days and transferred immediately to anaerobic conditions. Fresh samples were homogenized with sterile 1 × PBS (pH7.0) in an anaerobic glovebox (Electrotek, UK), which contained 10% hydrogen, 10% carbon dioxide and 80% nitrogen. The fecal suspension was diluted 10^−6^, 10^−7^, 10^−8^ and 10^−9^-fold, and plated on GAM medium (Nissui Pharmaceutical, Japan). Plates were incubated at 37°C for 3 days in an anaerobic glovebox. Single clones were picked and streaked until pure colonies were obtained on GAM medium. Full-length 16S rRNA gene sequencing was performed after amplification using primers (27 forward: 5’-AGAGTTTGATCCTGGCCTCAG-3’ and 1492 reverse: 5’-GGTTACCTTGTTACGACTT-3’). The isolates were stored at −80 °C in GAM broth containing 16% of glycerol until further use.

### Oxford nanopore sequencing

The strains 4-8-110 and 4-15-1 were recovered on GAM medium. Cells were harvested at the period of logarithmic growth. Genomic DNA was extracted using the Blood & Cell Culture DNA Midi Kit (Qiagen, Germany) following the manufacturer’s protocol. Libraries for whole-genome sequencing of the strains were constructed and sequenced on an ONT PromethION (Oxford Nanopore Technology, UK) at NextOmics (Wuhan, China). To correct sequencing errors, a library for second-generation sequencing was also constructed for each of the two strains and sequenced (2×100 bp) on a BGISEQ platform (BGI, China). Bioinformatic analyses of sequencing data were performed following Kolmogorov et al. (2019) and Hunt et al. (2015). In brief, after quality control, the sequence data were assembled with flye (Kolmogorov et al., 2019) with parameter: --nano-raw, and the assembled genomes were corrected by combining the Oxford Nanopore data with the second-generation sequencing data using pilon under default parameter. The encoded genes were predicted using prodial (parameter: -p none-g 11) (Hyatt et al., 2010).

### MAGs assembly

A total of 92 fecal samples from eight pig populations, four intestinal locations and different ages were used for metagenomic sequencing and construction of metagenome-assembled genomes (MAGs). Microbial DNA was extracted as described above. The libraries for metagenomic sequencing were constructed following the manufacturer’s instructions (Illumina, USA), with an insert size of 350 base pairs (bp) for each sample, and 2×150 bp paired-ends sequenced on a Novaseq 6000 platform. Raw sequencing data were filtered to remove adapter sequences and low-quality reads using fastp (v0.19.41) (Chen et al., 2018). Host genomic DNA sequences were filtered out using BWA (V.0.7.17) (Li & Durbin, 2010). The clean reads of each sample were assembled into contigs using MEGAHIT (v1.1.3) with the option ‘--min-count 2 --k-min 27 --k-max 87 --k-step 10 --min-contig-len 500’ (Li et al., 2016). Single-sample metagenomic binning was performed with two different binning algorithms ‘--metabat2 --maxbin2’ using the metaWRAP package (Uritskiy et al., 2018). The bins (metagenomic assembly genomes, MAGs) generated by the two binning algorithms were evaluated for quality and combined to form a MAG set using the bin_refinement module in metaWRAP. Metagenomic sequences were further assembled to optimized MAGs using the reassemble_bins module of metaSPAdes in the metaWRAP pipeline. CheckM was used to estimate the completeness and contamination of each MAG (Parks et al., 2015). The MAGs with completeness <50% and contamination >5% were filtered out. Non-redundant MAGs were generated by dRep (v2.3.2) at threshold of 99% average nucleotide identity (ANI) (Olm et al., 2017). The metagenomic sequencing data were submitted to the CNGB database and have accession number CNP0000824.

### Bioinformatic analyses of GalNAc catabolic pathway

Gene prediction in MAGs was carried out using the annotate_bins module in metaWRAP. The FASTA file of amino acid sequences translated from coding genes was used to perform KEGG annotation using Ghost KOALA tool (Kanehisa et al., 2016) on the KEGG website (https://www.kegg.jp/ghostkoala/). Taxonomic classification of MAGs was performed using PhyloPhlAn (v.0.99) (Segata et al., 2013). The graphs in Fig. 6B and 6C were generated using custom made perl and R scripts. Pathway and regulon scores were computed using a custom-made perl script. Both scores included (i) one point for import (having orthologues of either the four components of AgaPTS (agaE, agaF, agaV and agaW) and/or the three components of the TonB dependent transporter (omp, agaP and agaK) and /or the four components of the GnbPTS transporter (gnbA, gnbB, gnbC and gnbD), (ii) one point for GalNAc deacetylase activity (having an orthologue of agaA and/or nagA), (iii) one point for GalN deaminase/isomerase (having on orthologue of agaS), (iv) one point for tagatose-6-P kinase (having an orthologue of pfkA and/or lacC and/or fruK), and (v) one point for tagatose-1,6-PP aldolase (having an orthologue of gatY and/or gatZ and/or lacD and/or fba). For the pathway score the orthologues could be located anywhere in the MAG, for the regulon score they had to be located on the same sequence contig and in close proximity (2.5% of genome size) to the anchor gene agaS (Ravcheev & Tiele, 2017). For the top hits we manually checked whether proximity was confirmed either by the replication of the order in more than one MAG and/or by the colocalisation of the genes on one and the same sequence contig. The effect of MAG-type (OTU476-like, Erysipelotrichaceae and others), – completion, -contig, -genome size on pathway and regulon scores were estimated using the R lm function, and were highly significant. The p-values for the Erysipelotrichaceae versus Other contrast were directly obtained from the lm function. To (conservatively) estimate the p-value of the OTU476-like versus [Erysipelotrichaceae + Others] we generated score residuals corrected for completion, contig number and genome size and determined how many MAGs had scores as high or higher than the OTU476-like strains. The p-values reported in Fig. 6B correspond to the square of these proportions as there are two OTU476-like strains with same GalNAc cluster organization.

### Feeding experiment

To assess the effect of N-acetyl-galactosamine on the growth of bacterial strains 4-8-110 and 4-15-1 from Erysipelotrichaceae, a feeding experiment with α-N-acetyl-galactosamine was carried out in vitro. The OTU-476 like strains 4-8-110 and 4-15-1 were recovered and cultured on GAM broth medium. 0.005%, 0.05%, 0.1%, 0.2% and 0.5% of N-acetyl-galactosamine was added to the GAM broth medium, respectively. The GAM broth medium without N-acetyl-galactosamine was used as control. The two strains were inoculated in the above GAM broth medium and cultured at 37°C. The OD600 values of cultures at six different time points (0, 18, 25, 37 and 42h for 4-8-110, and 0, 11, 16, 18 and 24 h for 4-15-1) were measured using a UV Spectrophotometer (Yoke instrument, China). Student’s test was used to compare the abundance of the strains in GAM broth medium with and without N-acetyl-galactosamine.

### Profiling ABO gene expression level at various adult and embryo tissues

Total RNA was extracted using Trizol from 15 tissues (lung, hypophysis, skin, spinal cord, liver, spleen, muscle, hypothalamus, heart, blood, brain, cecum, stomach, duodenum and kidney) collected from an adult Bamaxiang sow and a Duroc pig embryo (day 75). RNA quality was monitored by agarose (1%) gel electrophoresis, and using the RNA Nano 6000 Assay Kit of the Bioanalyzer 2100 system (Agilent Technologies, CA, USA). RNA concentration was measured using Qubit^®^ RNA Assay Kit in Qubit^®^ 2.0 Flurometer (Life Technologies,CA, USA). 1-μg total RNA of each sample were used to constructed RNA sequencing libraries. Sequencing libraries were generated using TruSeq RNA Library Preparation Kit (Illumina, USA) following manufacturer’s recommendations and index codes were added to attribute sequences to each sample. Briefly, mRNA was purified from total RNA using poly-T oligo-attached magnetic beads. First strand cDNA was synthesized using random hexamer primer and M-MuLV Reverse Transcriptase (RNase H-). Second strand cDNA synthesis was subsequently performed using DNA Polymerase I and RNase H. Remaining overhangs were converted into blunt ends via exonuclease/polymerase activities. After adenylation of 3’ ends of DNA fragments, Illumina adaptors were ligated. In order to select cDNA fragments of preferentially 350~400 bp in length, the library fragments were purified with AMPure XP system (Beckman Coulter, Beverly, USA). PCR was performed with Phusion High-Fidelity DNA polymerase, Universal PCR primers and Index (X) Primer. PCR products were purified (AMPure XP system) and library quality was assessed on an Agilent Bioanalyzer 2100 system. Clustering of the index-coded samples was performed on a cBot Cluster Generation System using TruSeq PE Cluster Kit v3-cBot-HS (Illumina) according to the manufacturer’s instructions. Sequencing was performed on an Illumina Novaseq platform and 150 bp paired-end reads were generated. Reads were filtered obtained by removing adapter sequences, poly-N and low-quality reads. Cleaned reads were mapped to the complete ABO sequence from the Bamaxiang reference genome sequence using HISAT2 (Kim et al., 2019). Samtools (Li et al., 2009) was used to convert SAM format to BAM format. The read counts mapping to ABO (exon 1 to 7) were quantified for each sample using featureCounts (Liao et al., 2014). To adjust for the effect of sequencing depth, the expression abundance of ABO gene was normalized to Transcripts Per Million (TPM). Expression abundance of ABO was used to cluster and visualize the expression level of the 15 tissues from an adult Bamaxiang sow and a Duroc pig embryo via function dist(), hclust(), as.dendrogram() and set() implemented in R package stats and dendextend.

### Association analysis of ABO blood group with human gut microbiota

Human data used correspond to the previously described CEDAR cohort (Momozawa et al, 2018). It included 300 healthy individuals of European descent that were visiting the University Hospital (CHU) from the University of Liège as part of a national screening campaign for colon cancer. Blood samples and intestinal biopsies (ileum, colon and rectum) were collected with full consent. For microbiota analysis, DNA was extracted from biopsies using the QIAamp DNA Stool Mini Kit (QIAgen, Germany). Three 16S rRNA amplicons corresponding respectively to the V1-V2, V3-V4 and V5-V6 variable regions were generated in separate PCR reactions and subjected to paired-end (2×300bp) NGS sequencing on a MiSeq instrument (Illumina, USA) following Canver et al., 2015 at the GIGA genomics core facility. Reads were QV20 trimmed from the 3’ end, demultiplexed, primer sequences removed using the bbduk tool (BBMap – Bushnell B. – sourceforge.net/projects/bbmap/). Reads mapping to the human genome were eliminated using the BBTools suite [sourceforge.net/projects/bbmap/]. The corresponding pipeline was constructed using Snakemake (Köster & Rahmann, 2012). Further analyses were performed using QIIME 2 2018.11 (Bolyen et al., 2019). The paired end reads were denoised and joined using the DADA2 plugin (Callahan et al., 2016) using batch-specific trimming length parameters yielding 9.1±2.0K amplicon sequence variants (ASVs) per run for V1-V2, 4.5±1.6K for V3V4 and 6.8±0.67K for V5V6 amplicon. ASVs mapping to known contaminant taxa as well as ASVs with abundance negatively correlated with coverage depth were removed. Samples that more than 20% contaminant ASVs were eliminated from further analyses. ASVs were then clustered to 97% identity level OTUs using the DNACLUST program (Ghodsi et al., 2011). After OTU assignment, read counts were rarefied to 10,000 (V1-V2 and V5-V6) and 5,000 (V3-V4). As intestinal location only explored a minor proportion of the variance in OTU abundance (Shagam, in preparation), OTU abundances were averaged across locations. Local alignment identity of the detected ASVs with the OTU-476 and OTU-327 from the pig microbiome were measured using blastn (Altschul et al., 1990). The effect of ABO blood group on standardized abundances of individual OTUs was performed using a linear model (lm R function) including (i) ABO blood group (A, B, AB or O), (ii) secretor status, (iii) sex, (iv) smoking status, (v) age and (vi) BMI. Analyses were conducted separately for the different amplicons.

### Association analysis of 2.3-kb deletion of ABO gene with porcine complex traits

The associations between the 2.3Kb ABO deletion and 150 traits were calculated in the F6 and F7 populations based on a meta-analysis combining the effects. The observed P value for a trait was calculated by testing a weighted mean of Z scores from F6 and F7 generations as follows: Z = Z1W1+Z2W2/(W1+ W2), where Z1 = b1/SE1 and Z2 = b2/SE2, W1 = 1/(SE1)2 and W2 = 1/(SE2)2, where the subscripts 1 and 2 denote F6 and F7 generations, respectively; b1, b2, SE1 and SE2 were effects and standard errors of ABO locus on a given trait estimated from a linear mixed model, which accounted for population structure using a genomic relationship matrix derived from whole genome marker genotypes. A total of 250 and 254 traits were tested in F6 and F7 generation, while 150 traits that were shared in F6 and F7 generations were used for meta-analysis.

## Author contributions

HY analyzed the 16S rRNA sequencing data, performed the GWAS, meta-analyses and local association analyses, computed heritabilities of individual taxa, contributed to ABO genotyping and analyzed the effect of the 2.3 Kb deletion on taxa abundance. JW analyzed the composition of the microbiome including PCoA analyses, computation of *β* – and *α* – diversity, computed correlations between kinship and microbiome dissimilarities and their significance, isolated the OTU476-like strains, performed the GalNAc feeding experiments, measured the concentrations of GalNAC in cecal lumen, analyzed the GalNAc import and utilization pathway in the MAGs, and contributed to ABO genotyping. XH participated in 16S rRNA sequencing (F6) and GWAS (F6). YZ performed metagenome sequencing analysis, analyzed the GalNAc import and utilization pathway in the MAGs, analyzed the RNA seq data from cecum samples, and contributed to ABO genotyping. YZ participated in the preparation of the genotype data from whole genome sequence information, participated in the computation of the genomic contribution of the different breeds in the F6 and F7 generation and the definition of expected mapping resolution, performed LD analyses, performed eQTL analysis for the ABO gene, participated in the characterization and sequence analysis of the ABO gene including definition of the 2.3 Kb deletion, and participated in the balancing selection and trans-species polymorphism analyses. ML assisted with the isolation of the OTU476-like strains, the GalNAc feeding experiments, and genotyping of the ABO gene. QL assisted with measuring the concentrations of GalNAc in cecal lumen. SK, MH, HF, SF, XX, HJ, SC and JG assisted with the experiments. XT determined the expression profiles of ABO gene in different tissues of adult and fteus pigs. ZZ, ZW, HG and YH assisted with the preparation of genotype data from whole-genome sequence data and conducted the analysis of the Nanopore data of the ABO region. JM assisted with the construction of the mosaic population. HA assisted with the bioinformatic analysis of the ABO region, the de novo assembly of the A allele, and the evolutionary analysis of the ABO alleles. LS analyzed the effect of ABO genotype on intestinal microbiota composition in humans. WC assisted in the analysis of the sequence data for the trans-species polymorphisms analysis. CaCh supervised the characterization of the ABO gene and the 2.3 Kb deletion and the corresponding haplotype structure in the F0, F6 and F7 population and for the trans-species polymorphism. BY prepared the genotype data of whole-genome variants, assisted with the raising of swine heterogeneous stock, and participated in the computation of the genomic contribution of the different breeds in the F6 and F7 generation and the definition of expected mapping resolution. MG supervised the bioinformatic and statistical analyses, performed bioinformatic and statistical analyses, and wrote the paper. CC codesigned the study, supervised experiments, supervised bioinformatic and statistical analyses of gut microbiome and wrote the paper. LH created the swine heterogeneous stock, designed the study, directed the project, supervised the experiments and analyses, and wrote the paper.

## Data accessibility

The 16S rRNA sequencing data and the genotype data of the F0, F6 and F7 were submitted to the CNGB database and have accession number CNP0001069. The metagenomic sequence data were submitted to the CNGB database and have accession number CNP0000824.

## Acknowledgements

We are very grateful to Yuyong He, Shijun Xiao, Wanbo Li, Yuanmei Guo and Yuyun Xing for their great assistance in the construction of the experimental mosaic pig populations, and to Ying Su and Jing Li for their preparation of reagents and management of samples. We are also grateful to Yukihide Momozawa, Rob Mariman, Myriam Mni, Latifa Karim and Manon Dekkers for generating the CEDAR-1 16S rRNA data. Lusheng Huang is supported by The National Natural Science Foundation of China (31790410) and National pig industry technology system (CARS-35). We thank the long-term projects support from Jiangxi department for education, the Ministry of Science and Technology of P.P. China, the Ministry of Agriculture and Rural Affairs of P. R. China, and Jiangxi department of Science and Technology for the swine heterogeneous stock. Congying Chen was supported by the National Natural Science Foundation of China (31772579). Hui Yang is supported by National Postdoctoral Program for Innovative Talent (No. BX201700102). Lev Shagam is supported by the FNRS IBD-GI-Seq project to Souad Rahmouni. Michel Georges is supported by the Chinese Thousand Talents Program and the Belgian EOS “Miquant” project. Carole Charlier is Senior Research Associate from the FNRS.

## References

Ai, H., et al. Adaptation and possible ancient interspecies introgression in pigs identified by whole-genome sequencing. Nat Genet 47, 217–225 (2015).

Aleman, F.D.D., Valenzano, D.R. Microbiome evolution during hots aging. PLoS Pathog 15, e1007727 (2019).

Altschul, S.F., Gish, W., Miller, W., Myers, E.W., Lipman, D.J. Basic alocal alignment search tool. J Mol Biol 215, 403–410 (1990).

Aulchenko, Y. S., Ripke, S., Isaacs, A. & van Duijn, C. M. GenABEL: an R library for genome-wide association analysis. Bioinformatics 23, 1294–1296 (2007).

Bidart, G.N., Rodriguez-Diaz, J., Monedoro, V., Yebra, M.J A unique gene cluster for the utilization of the mucosal and human milk-associated glycans galacto-N-biose and lacto-N-biose in *Lactobacillus casei*. Mol Microbiol 93, 521–538 (2014).

Blancher, A. Evolution of the ABO supergene family. ISBT Science Series 8, 201–206 (2013).

Blekhman, R. et al. Host genetic variation impacts microbiome compoistion across human body sites. Genome Biol 16, 191 (2015).

Bolger, A.M., Lohse, M., Usadel, B. Trimmomatic: a flexible trimmer for Illumina sequence data. Bioinformatics 30, 2114–2120 (2014).

Bolyen, E. et al. Reproducible, interactive, scalable and extensible microbiome data science using QIIME2. Nat Biotechnol 37, 852–857 (2019).

Bonder, M.J. et al. The effect of host genetics on the gut microbiome. Nat Genet 48, 1407–1412 (2016).

Boren, T. et al. Attachment of Helicobacter pylori to human gastric epithelium mediated by blood group antigens. Science 262, 1892–1895 (1993).

Boyle, E.A., Li, Y.I., Pritchard, J.K. An expanded view of complex traits: from polygenic to omnigenic. Cell 169, 1177–1186 (2017).

Bray, J.R. and Curtis, J.T. An ordination of uplnad forest communities of southern Wisconsin. Ecologial Monographs 27, 325–349 (1957).

Brinkkötter, A.B., Klöss, H., Alpert, C.-A., Lengeler, J.W. Pathways for the utilization of N-acetyl-galactosamine and galactosamine in Escherichia coli. Mol Microbiol 37, 125–135 (2000).

Browning, S. R. & Browning, B. L. Rapid and accurate haplotype phasing and missing-data inference for whole-genome association studies by use of localized haplotype clustering. Am J Hum Genet 81, 1084–1097 (2007).

Callahan, B.J., McMurdie, P.J., Rosen, M.J., Han, A.W., Johnson, A.J.A., Holmes, S.P. DADA2: high-resolution sample inference from Illumina amplicon data. Nat Methods 13, 581–583 (2016).

Camus, D., Bina, J.C., Carlier, Y., Santoro, F. ABO blood groups and clinical forms of schistosomiasis mansoni. Trans R Soc Trop Med Hyg 71, 182 (1977).

Chang, C. C. et al. Second-generation PLINK: rising to the challenge of larger and richer datasets. Gigascience 4, 7 (2015).

Charlier, C., Li, W., Harland, C., Littlejohn, M., Coppieters, W., Creagh, F., Davis, S., Druet, T., Faux, P., Guillaume, F., Karim, L., Keehan, M., Kadri, N.K., Tamma, N., Spelman, R., Georges, M. NGS-based reverse genetic screen for common embryonic lethal mutations compromising fertility in livestock. Genome Res 26, 133–1341 (2016).

Chaudhuri, A., De S. Cholera and blood groups. Lancet ii, 404 (1977).

Cheema, M.U., Pluznick, J.L. Gut Microbiota Plays a Central Role to Modulate the Plasma and Fecal Metabolomes in Response to Angiotensin II. Hypertension 74, 184–193 (2019).

Chen, S., Zhou, Y., Chen, Y. & Gu, J. fastp: an ultra-fast all-in-one FASTQ preprocessor. Bioinformatics 34, i884–i890 (2018).

Chen, Y. et al. ABO blood group and susceptibility to severe acute respiratory syndrome. JAMA 293, 1450–1451 (2005).

Choi, M.K., Le, M.T., Cho, H., Yum, J., Kang, M., Song, H., Kim, J.H., Chung, H.J., Hong, K., Park, C. Determination of complete sequence information of the human ABO blood group orthologous gene in pigs and breed differences in blood type frequencies. Gene 640, 1–5 (2018).

Cooling, L. Blood groups in infection and host susceptibility. Clin Microbiol Rev 28, 801–870 (2015).

Coppieters, W., Karim, L., Georges, M. SNP-based deconvolution of biological mixtures: application to the detection of cows with subclinical mastitis by whole genome sequencing of tank milk. Genome Res, in the press (2020).

Davenport, E.R, Goodrich, J.K., Cell, J.T., Spector, T., Ley, R.E., Clark, A.G. ABO antigen and secretor statuses are not associated with gut microbiota composition in 1,500 twins. BMC Genomics 17, 941–955 (2016).

DePristo, M., Banks, E., Poplin, R., Garimella, K., Maguire, J., Hartl, C., Philippakis, A., del Angel, G., Rivas, M.A., Hanna, M., McKenna, A., Fennell, T., Kernytsky, A., Sivachenko, A., Cibulskis, K., Gabriel, S., Altshuler, D., Daly, M. A framework for variation discovery and genotyping using next-generation DNA sequencing data. Nat Genet 43, 491–498 (2011).

DeSantis, T.Z., Hugenholtz, P., Larsen, N., Rojas, M., Brodie, E.L., Keller, K., Huber, T., Dalevi, D., Hu, P., Andersen, G.L. Greengenes, a chimera-checked 16S rRNA gene database and workbench compatible with ARB. Appl Environ Microbiol 72, 5069–5072 (2006).

Dixon P. VEGAN, a package of R functions for community ecology. J Veg Sci 14, 927–930 (2003).

Dobin, A., Davis, C.A., Schlesinger, F., Drenkow, J., Zaleski, C., Jha, S., Batut, P., Chaisson, M., Gingeras, T.R. STAR: ultrafast universal RNA-Seq aligner. Bioinformatics 29, 15–21 (2013).

Donaldson, G.P., Lee, S.M., Mazmanian, S.K. Gut biogeography of the bacterial microbiota. Nat Rev Microbiol 14, 20–32 (2016).

Edgar, R.C. Search and clustering orders of magnitude faster than BLAST. Bioinformatics 26, 2460–2461 (2010).

Ellinghaus, D. et al. The ABO blood group locus and a chromosome 3 gene cluster associate with SARS-CoV-2 respiratory failure in an Italian-Spanish genome-wide association analysis. medRxiv, https://www.medrxiv.org/content/10.1101/2020.05.31.20114991v1 (2020).

Falconer, D.S., Mackay, T.F.C. Introduction to quantitative genetics. 4^th^ Edition. Pearson education Limited. (1996).

Frantz, L.A.F. et al. Evidence of long-term gene flow and selection during domestication from analyses of Eurasian wild and domestic pig genomes. Nat Genet 47, 1141–1148 (2015).

Georges, M., Charlier, C., Hayes, B. Harnessing genomic information for livestock improvement. Nat Rev Genet 20, 135–156 (2019).

Gerlades, A. et al. Inferring the history of speciation in house mice from autosomal, X-linked, Y-linked and mitochondrial genes. Mol Ecol 17, 5349–5363 (2008).

Ghodsi, M., Liu, B., Pop, M. DNACLUST: accurate and efficient clsutering of phylogenetic marker genes. BMC Bioinformatics 12, 271 (2011).

Goodrich, J.K., Waters, J.L., Poole, A.C., Sutter, J.L., Koren, O., Blekhman, R., Beaumont, M., Van Treuren, W., Knight, R., Bell, J.T., Spector, T.D., Clark, A.G., Ley, R.E. Human genetics shape the gut microbiome. Cell 159, 789–799 (2014).

Goodrich, J.K., Davenport, E.R., Beaumont, M., Jackson, M.A., Knight, R., Ober, C., Spector, T.D., Bell, J.T., Clark, A.G., Ley, R.E. Genetic Determinants of the Gut Microbiome in UK Twins. Cell Host Microbe 19, 731–743 (2016).

Groenen, M.A.M. A decade of pig genome sequencing: windo on pig domestication and evolution. Genet Sel Evol 48, 23–32 (2016).

Hanson, M.E.B. et al. Population structure of human gut bacteria in a diverse cohort from rural Tanzania and Botswana. Genome Biology 20, 16 (2019).

Harris, R. S. Improved pairwise alignment of genomic DNA. Doctor thesis, The Pennsylvania State University (2007).

Haseman, J.K. and Elston, R.C. The investigation of linkage between a quantitative trait and a marker locus. Behav Genet 2, 3–19 (1972).

Hughes, D.A., Bacigalupe, R., Wang, J., Ruhlemann, M.C., Tito, R.Y., Falony, G., Joossens, M., Vieira-Silva, S., Henckaerts, L., Rymenans, L., et al. Genome-wide associations of human gut microbiome variation and implications for causal inference analyses. Nat Microbiol, Online ahead of print (2020). 10.1038/s41564-020-0743-8

Hunt, M. et al. Circlator: automated circularization of genome assemblies using long sequencing reads. Genome Biol 16, 294 (2015).

Hu, Z., Patel, I.R. & Mukherjee, A. Genetic analysis of the roles of agaA, agaI, and agaS genes in the N-acetyl-D-galactosamine and D-galactosamine catabolic pathways in Escherichia coli strains O157:H7 and C. BMC Microbiol 13, 94 (2013).

Huang, H. et al. Fine-mapping inflammatory bowel disease loci to single-variant resolution. Nature 547, 173–178.

Hyatt, D., et al. Prodigal: prokaryotic gene recognition and translation initiation site identification. BMC Bioinformatics 11, 119 (2010).

Kadri, N.K. et al. A 660-Kb deletion with antagonistic effects on fertility and milk production segregates at high frequency in Nordic red cattle: additional evidence for the common occurrence of balancing selection in livestock. PLoS Genet 10, e1004049 (2014)

Kanehisa, M., Sato, Y., Morishima, K. BlastKOALA and GhostKOALA: KEGG tools for functional characterization of genome and metagenome sequences. J Mol Biol 428, 726–731 (2016).

Kelly, R.J., Rouquier, S., Giorgi, D., Lennon, G.G., Lowe, S.B. Sequence and expression of a candidate for the human Secretor blood group *α*(1,2) fucosyltransferase gene (*FUT2*). J Biol Chem 270, 4640–4649 (1995)

Kittelman, S., et al. Two different bacterial community types are linked with the low-methane emission trait in sheep. PLoS ONE 9, e103171 (2014).

Kolmogorov, M., Yuan, J., Lin, Y. & Pevzner, P. A. Assembly of long, error-prone reads using repeat graphs. Nat Biotechnol 37, 540–546 (2019).

Koonin, E.V. Evolution of genome architecture. Int J Biochem Cell Biol 41, 298–306 (2009)

Köster, J. and Rahmann, S. Snakemake: a scalable bioinformatics workflow engine. Bioinformatics 28, 2520–2522 (2012).

Kundu, P., Blacher, E., Elinav, E., Pettersson, S. Our gut microbiome: the evolving inner self. Cell 171, 1481–1493 (2017).

Lawrence, J. Selfish operons: the evolutionary impact of gene clustering in prokrayotes and eukaryotes. Curr Opin Genet Dev 9, 642–648 (1999).

Leyn, S.A., Gao, F., Yang, C., Rodionov, D.A. N-acetylgalactosamine utilization pathway and regulon in proteobacteria. J Biol Chem 287, 28047–28056 (2012).

Li, H., Handsaker, B., Wysoker, A., Fennell, T., Ruan, J., Homer, N., Marth, G., Abecassis, G., Durbin, R. The sequence alignment/map format and SAMtools. Bioinformatics 25, 2078–2079 (2009).

Li, H., Durbin, R. Fast and accurate long-read alignment with Burrows–Wheeler transform. Bioinformatics 26, 589–595 (2010).

Li, H. Aligning sequence reads, clone sequences and assembly contigs with BWA-MEM. arXiv:1303.3997 (2013).

Li, D., et al. MEGAHIT v1.0: A fast and scalable metagenome assembler driven by advanced methodologies and community practices. Methods 102, 3–11 (2016).

Li, H. Minimap2: pairwise alignment for nucleotide sequences. Bioinformatics 34, 3094–3100 (2018).

Liao, Y., Smyth, G.K., Shi, W. featureCounts: an efficient general-purpose program for assigning sequence reads to genomic features. Bioinformatics 30, 923–930 (2014).

Lindesmith, L. et al. Human susceptibility and resistance to Norwalk virus infection. Nat Med 9, 548–553 (2003).

Magoc, T. & Salzberg, S. L. FLASH: fast length adjustment of short reads to improve genome assemblies. Bioinformatics 27, 2957–2963 (2011).

Mahowald, M.A., Rey, F.E., Seedorf, H., Turnbaugh, P.J., Fulton, R.S., Wollam, A. et al. Characterizing a model human gut microbiota composed of members of its two dominant bacterial phyla. Proc Natl Acad Sci USA 106, 5859–5864 (2009).

Makivuokko, H., Lahtinen, S.J., Wacklin, P., Tuovinen, E., Tenkanen, H., Nikkila, J., Bjorklund, M., Aranko, K., Ouwenhand, A.C., Matto, J. Association between the ABO blood group and the human intestinal microbiota composition. BMC Microbiol 12, 94 (2012).

Malmuthuge, N., Griebel, P.J., Guan, L.L. Taxonomic identification of commensal bacteria associated with the mucosa and digesta throughout the gastrointestinal tracts of preweaned calves. Appl Environ Microbiol 80, 2021–2028 (2014).

Momozawa, Y. et al. IBD risk loci are enriched in multigenic regulatory modules encompassing putative causative genes. Nat Commun 9, 2427 (2018).

Ndamba, J., Gomo, E., Nyazema, N., Makaza, N., Kaondera, K.C. Schistosomiasis infection in relation to the ABO blood groups among school children in Zimbabwe. Acta Trop 65, 181–190 (1997).

Nei, M. F-statistics and analysis of gene diversity in subdivided populations. Ann Hum Genet 41, 225–233 (1977).

O’Hara, E., Neves, A.L.A., Song, Y., Guan, L.L. The role of the gut microbiome in cattle production and health: driver or passenger? Annu Rev Anim Biosci 8, 199–220 (2020).

Olm, M.R., Brown, C.T., Brooks, B. & Banfield, J.F. dRep: a tool for fast and accurate genomic comparisons that enables improved genome recovery from metagenomes through de-replication. ISME J 11, 2864–2868 (2017).

Parks, D.H., Imelfort, M., Skennerton, C.T., Hugenholtz, P. & Tyson, G.W. CheckM: assessing the quality of microbial genomes recovered from isolates, single cells, and metagenomes. Genome Res 25, 1043–1055 (2015).

Patterson, N., et al. Genetic evidence for complex speciation of humans and chimpanzees. Nature 441, 1103–1108 (2006).

Pereira, F.E.L., Bortolini, E.R., Carneiro, J.L.A., da Silva, C.R.M., Neves, R.C. A,B,O blood groups and hepatosplenic form of schistosomiasis mansoni (Symmer’s fibrosis). Trans R Soc Trop Med Hyg 71, 182 (1977).

Polderman, T.J.C. et al. Meta-analysis of the heritability of human traits based on 50 years of twin studies. Nat Genet 47, 702–709 (2015).

Polubriaginof, F.C.G. et al. Disease heritability inferred from familial relationships reported in medical records. Cell 173, 1692–1704 (2018).

Quast, C., Pruesse, E., Yilmaz, P., Gerken, J., Schweer, T., Yarza, P., Peplies, J., Glöckner, F.O. The SILVA ribosomal RNA gene database project: improved data processing and web-based tools. Nucleic Acids Res 41, D590–D596 (2013).

Radjabzadeh D, Boer CG, Beth SA, van der Wal P, Kiefte-De Jong JC, Jansen MAE, Konstantinov SR, Peppelenbosch MP, Hays JP, Jaddoe VWV, Ikram MA, Rivadeneira F, van Meurs JBJ, Uitterlinden AG, Medina-Gomez C, Moll HA, Kraaij R. Diversity, compositional and functional differences between gut microbiota of children and adults. Sci Rep 10, 1040 (2020).

Ravcheev, D.A., Thiele, I. Comparative genomic analysis of the human gut microbiome reveals a broad distribution of metabolic pathways for the degradation of host-synthesized mucin glycans and utilization of mucin-derived monosaccharides. Front Genet 8, 111 (2017).

Rimmer, A. et al. Integrating mapping-, assembly- and haplotype-based approaches for calling variants in clinical sequencing applications. Nat Genet 46, 912–918 (2014).

Rodionov, D.A., Yang, C., Li, X., Rodionova, I.A., Wang, Y., Obraztsova, A.Y., Zagnitiko, O.P. Overbeek, R., Romine, M.F., Reed, S., Frederickson, J.F., Nealson, K.H., Osterman, A.L. Genomic encyclopedia of sugar utiulization pathways in the Shewanella genus. BMC Genomics 11, 494 (2010).

Rognes, T., Flouri, T., Nichols, B., Quince, C. & Mahe, F. VSEARCH: a versatile open source tool for metagenomics. PeerJ 4, e2584 (2016).

Ross, E.M., Moate, P.J., Marett, L.C., Cocks, B.G., Hayes, B.J. Metagenomic predictions: from microbiome to complex health and environmental phenotypes in human and cattle. PLOS ONE 8, e73056 (2013).

Rothschild, D., et al. Environment dominates over host genetics in shaping human gut_microbiota. Nature 555, 210–215 (2018).

Rowe, J.A. et al. Blood group O protects against severe Plasmodium falciparum malaria through the mechanism of reduced rosetting. Proc Natl Acad Sci USA 104, 17471–17476 (2007).

Sankararaman, S., Mallick, S., Dannemann, M., Prüfer, K., Kelso, J., Pääbo, S., Patterson, N., Reich, D. The genomic landscape of Neanderthal ancestry in present-day humans. Nature 507, 354–357 (2014).

Schlamp, F., Zhang, D.Y., Cosgrove, E., Simecek, P., Edwards, M., Goodrich, J.K., Ley, R.E., Churchill, G.A., Clark, A.G. High-resolution QTL mapping with diversity outbred mice identifies genetic variants that impact gut microbiome composition. bioRxiv https://doi.org/10.1101/722744 (2019).

Schloss, P. D. et al. Introducing mothur: open-source, platform-independent, community-supported software for describing and comparing microbial communities. Applied and environmental microbiology 75, 7537–7541 (2009).

Schmidt, T.S.B., Raes, J., Bork, P. The human gut microbiome: from association to modulation. Cell 172, 1198–1215 (2018).

Segata, N., Bornigen, D., Morgan, X.C. & Huttenhower, C. PhyloPhlAn is a new method for improved phylogenetic and taxonomic placement of microbes. Nat Commun 4, 2304 (2013).

Ségurel, L., Thompson, E.E., Flutre, T., Lovstad, J., Venkat, A., Margulis, S.W., Moyse, J., Ross, S., Gamble, K., Sella, G., Ober, C., Przeworski, M. The ABO blood group is a trans-species poilymorphism in primates. Proc Natl Acad Sci USA 109, 18493–18498 (2012).

Srivastava, A., et al. Genomes of the mouse collaborative cross. Genetics 206, 537–556 (2017).

The 1000 Genomes project Consortium. A map of human genome variation from population-scale sequencing. Nature 467, 1061–1073 (2010)

Turpin, W. et al. Association of host genome with intestinal microbial composition in a large healthy cohort. Nature genetics 48, 1413–1417 (2016).

Uritskiy, G.V., DiRuggiero, J. & Taylor, J. MetaWRAP-a flexible pipeline for genome-resolved metagenomic data analysis. Microbiome 6, 158 (2018).

Vaser, R., Sovic, I., Nagarajan, N. & Sikic, M. Fast and accurate de novo genome assembly from long uncorrected reads. Genome Res 27, 737–746 (2017).

Van der Auwera, G. A. et al. From FastQ data to high confidence variant calls: the Genome Analysis Toolkit best practices pipeline. Curr Protoc Bioinformatics 43, 11.10.11–11.10.33 (2013).

Visscher, P.O., et al. Assumption-free estimation of heritability from genome-wide identity-by-descent sharing between full-sibs. PLoS Genet 2, e41 (2006).

Walker, B. J. et al. Pilon: an integrated tool for comprehensive microbial variant detection and genome assembly improvement. PLoS One 9, e112963 (2014).

Walter J., Armet, A.M., Brett Finlay, B., Shanahan, F. Establishing or exaggerating causality for the gut microbiome: lessons from human microbiota-associated rodents. Cell 180, 221–231 (2020).

Wang, Q., Garrity, G. M., Tiedje, J. M. & Cole, J. R. Naive Bayesian classifier for rapid assignment of rRNA sequences into the new bacterial taxonomy. Applied and environmental microbiology 73, 5261–5267 (2007).

Wang, M., Pryce, J.E., Savin, K., Hayes, B.J. Prediction of residual feed intake from genome and metagenome profiles in first lactation Holstein-Friesian dairy cattle. Proc. Assoc. Adv. Breed. Genet. 21, 89–92 (2015).

Wang, S., Cuesta-Seijo, J.A., Striebeck, A., Lafont, B., Palcic, M.M., Vidal, S. Design of glycosyl transferase inhibitors: serine analogues as pyrophosphate surrogates? ChemPlusChem 80, 1525–1532 (2015).

Wang, J. et al. Genome-wide association analysis identifies variation in vitamin D receptor and other host factors influencing the gut microbiota. Nat Genet 48, 1396–1406 (2016).

Warr, A., Affara, N., Aken, B., Beiki, H., Bickhart, D. M., Billis, K., Chow, W., Eory, L., Finlayson, F. A., Flicek, P., Girón, C. G., Griffin, D. K., Hall, R., Hannum, G., Hourlier, T., Howe, K., Hume, D. A., Izuogu, O., Kim, K., Koren, S., Liu, H., Manchanda, N., Martin, F. J., Nonneman, D. J., O’Connor, R. E., Phillippy, A. M., Rohrer, G. A., Rosen, B. D., Rund, L. A., Sargent, C. A., Schook, L. B., Schroeder, S. G., Schwartz, A. S., Skinner, B. M., Talbot, R., Tseng, E., Tuggle, C. K., Watson, M., Smith, T. P. L. & Archibald, A. L. An improved pig reference genome sequence to enable pig genetics and genomics research. bioRxiv, doi: https://doi.org/10.1101/668921.

Weir, B.S., Cockerham, C.C. Estimating F-statistics for the analysis of population structure. Evolution 38, 1358–1370 (1984).

Willer, C.J., Li, Y., Abecasis, G.R. METLA: fast and efficient meta-analysis of genomewide association scans. Bioinformatics 26, 2190–2191 (2010).

Yang, J., Lee, S. H., Goddard, M. E. & Visscher, P. M. GCTA: a tool for genome-wide complex trait analysis. Am J Hum Genet 88, 76–82 (2011).

Yatsunenko, T. et al. Human gut microbiome viewed across age and geography. Nature 486, 222–227 (2012).

Yengo, L. et al. Meta-analysis of genome-wide association studies for height and body mass index in ~700,000 individuals of European ancestry. Hum Mol Genet 27, 3641–3649 (2018).

Yu, N., Zhao, Z., Fu, Y.-X., Sambuughin, N., Ramsay, M., Jenkins, T., Leskinen, E., Patthy, L., Jorde, L.B., Kuromori, T., Li, W.-H. Global patterns of human DNA sequence variation in a 10-kb region on chromosome 1. Mol. Biol. Evol. 18, 214–222 (2001).

Zhang, H., Ravcheev, D.A., Hu, D., Zhang, F., Gong, X., Hao, L., Cao, M., Rodionov, D.A., Wang, C., Feng, Y. Tow novel regulators of N-acetyl-galactosamine utilization pathway and distinct roles in bacterial infections. Microbiol Open 4, 983–1000 (2015).

Zhou, X. & Stephens, M. Genome-wide efficient mixed-model analysis for association studies. Nat Genet 44, 821–824 (2012).

